# Polycomb group protein CBX7 represses cardiomyocyte proliferation via modulation of the TARDBP/Rbm38 axis

**DOI:** 10.1101/2021.09.08.459291

**Authors:** Kyu-Won Cho, Mark Andrade, Seongho Bae, Sangsung Kim, Jin Eyun Kim, Er Yearn Jang, Ahsan Husain, Roy L. Sutliff, John W. Calvert, Changwon Park, Young-sup Yoon

## Abstract

Cardiomyocyte (CM) proliferation notably decreases during the perinatal period. At present, regulatory mechanisms for this loss of proliferative capacity is poorly understood. CBX7, a polycomb group (PcG) protein, regulates the cell cycle but its role in CM proliferation is unknown. Here, we report that CBX7 inhibits proliferation of perinatal CMs by controlling TARDBP/Rbm38 pathway. Gene expression profiling demonstrated that CBX7 expression in the heart was low during the prenatal period, abruptly increased during the perinatal period, and sustained constantly throughout the adulthood. CBX7, when overexpressed via adenoviral transduction in neonatal CMs, reduced proliferation and promoted multinucleation of the CMs. Mutant mice carrying targeted inhibition of CBX7 in CMs exhibited cardiomegaly with increased proliferation of CMs at postnatal stages. Mechanistically, CBX7 interacted with TAR DNA-binding protein 43 (TARDBP) and positively regulated its downstream target, RNA Binding Motif Protein 38 (RBM38). Rbm38 was upregulated in the postnatal hearts and overexpression of RBM38 reduced proliferation of neonatal CMs. Together, this study provides a novel insight into the role of CBX7 in regulation of CM proliferation during the perinatal period.

## Introduction

Heart failure is one of the leading causes of mortality and morbidity in the world, and no effective treatment is available due to the limited regenerative capacity of the adult heart after injury^1–3^. Attempts have been made towards inducing cardiomyocyte (CM) proliferation, since it is considered one of the key processes for heart regeneration. This concept has been supported by a series of studies using genetic model organisms including zebrafish and neonatal mouse, in which the regenerative response accompanied robust division of pre-existing CMs^4–6^. Similarly, pre-existing CMs are the primary source of newly generated CMs in the adult mammalian heart during aging and after injury despite at a low frequency^7^. The CM renewal declines beginning at the perinatal stage^8, 9^; however, it is yet unclear why and how mammalian CMs lose proliferative capacity after birth.

It has been known that cell proliferation is controlled by Polycomb group (PcG) proteins via two mechansms^10, 11^. First, they are present in the nuclei and epigenetically control transcription of cell cycle-regulatory genes such as CDKN2A^12^, Cyclin A^13^, PTEN^14^, and c-MYC^15^. The canonical mechanism of PcG-mediated epigenetic silencing involves coordinated actions of two major types of Polycomb repressive complex (PRC), PRC 1 and 2^16^. PRC2, consisting of EZH1/2, SUZ12, EED, and RBBP4/7, initiates the repression process by tri-methylation of histone 3 tail (H3K27me3)^17^. PRC1, composed of RING1A/B, PCGF1-6, CBX family, PHC1-3, and SCMH1/2, is then recruited and stabilizes this silencing process via mono-ubiquitination of H2A tail (H2Aub)^18^. Finally, H2Aub serves as a binding site for PRC2, which further propagates the H3K27me3 repressive histone mark on H2Aub nucleosomes, generating a positive feedback loop^19^. While much less well known, PcG proteins are also present in the cytoplasm and control cell proliferation. EZH2 forms cytosolic PRC2 together with EED and SUZ12, and controls receptor-mediated cell proliferation in fibroblasts and T cells via its methyltransferase activity^20^.

A few PcG proteins were reported to regulate CM proliferation through epigenetic regulation of transcriptional programs. Deletion of EZH2 in cardiac progenitors at an early embryonic stage reduced CM proliferation and induced cardiac defects, leading to perinatal lethality^21^. The mechanism was proposed that EZH2 induces H3K27 trimethylation (H3K27Me3) at the loci of cyclin-dependent kinase inhibitors such as Ink4a/b in fetal CMs. CM-specific deletion of EED1 also reduced CM proliferation, causing thinning of myocardial walls and embryonic lethality although the mechanism has not been elucidated^21, 22^. Studies showed that JMJ repressed CM proliferation by epigenetically downregulating cyclin D1 and by directly interacting with retinoblastoma protein^23, 24^. In addition, deletion of JMJ resulted in congenital cardiac defects such as excessive trabeculation, double outlet right ventricle (DORV), and ventricular septal defect (VSD)^25^. Phc1 knockout mice also exhibited cardiac defects including looping defect and VSD^26^. These reports suggest that PcG proteins play critical roles in CM proliferation and cardiac development.

CBX7 is one of the PRC1 subunits and has been suggested to regulate cell proliferation mostly in cancer cells^27–35^. However, studies reported opposite functions of CBX7 in cellular proliferation as an oncogene or a tumor suppressor. These divergent observations on CBX7 function suggest that the role of CBX7 could be tissue-specific and context-specific^36^. CBX7 acts as a reader for H3K27me3 and mediates stabilization of heterochromatin, leading to transcriptional repression of target genes^18, 37^. However, we discovered that CBX7 exists as two different alternative splicing isoforms: 36 kDa and 22 kDa proteins (p36^CBX7^ and p22^CBX7^, respectively). These two CBX7 isoforms are found in mammals and exhibit distinct characteristics. p36^CBX7^ is localized to the nucleus and endogenously expressed in proliferating cells whereas p22^CBX7^ is localized to the cytoplasm and expressed under serum deprivation, inhibiting cell proliferation when overexpressed^38^. Thus, their exclusive function as an epigenetic regulator is questionable due to the presence of its cytoplasmic isoform. More importantly, the role of CBX7 in CM proliferation and cardiac development has been undescribed.

In this study, we demonstrated that CBX7 is mainly expressed as the p22^CBX7^ cytoplasmic isoform in CMs and represses proliferation of CMs. During the transition from the prenatal to the postnatal stage, CBX7 became abruptly upregulated in the heart, which was sustained into and during adulthood. CBX7 overexpression reduced proliferation of neonatal CMs and CM-specific inhibition of CBX7 in conditional knock-out mice resulted in increased CM proliferation *in vivo*. Mechanistically, CBX7 interacted with TARDBP, a versatile mitosis regulator, and upregulated its downstream target, Rbm38, an RNA binding protein regulating cellular proliferation. Together, this study uncovers a novel mechanism in which a PcG protein CBX7 present in cytoplasm can function as a molecular switch to restrict proliferation of CMs during the perinatal period.

## Results

### Temporal Changes of Polycomb group proteins in the heart

To investigate whether PcG proteins are involved in the regulation of CM proliferation, we examined expression patterns of 18 PcG genes that are reported to control the cell cycle^11^ (Cbx2, Cbx4, Cbx6, Cbx7, Cbx8, Pcgf1-6, Phc1, Ring1-2, Scmh1, Ezh2, Suz12 and Yy1) in the fetal (E17.5), neonatal (P0) and adult (5-month-old) hearts by quantitative RT-PCR (qRT-PCR). Among them, 9 genes including Cbx7, Cbx4, and Pcgf1 were upregulated in the neonatal heart compared to the fetal heart (**Figure 1A**). When comparing the gene expression between adult heart and neonatal heart, only Cbx7 was upregulated in the adult heart whereas 11 genes such as Ezh2, Pcgf2, and Ring2 became downregulated (**Figure 1A**). When comparing adult and fetal hearts, only Cbx7 was upregulated and 6 genes (Pcgf2, Ezh2, Ring2, Pcgf3, Phc1, and Yy1) were downregulated in adult hearts (**Figure 1A**). These results indicate that Cbx7 is the only gene significantly upregulated during the transitions from both fetal-to-neonatal and neonatal-to-adult stages with a total increase of 15.4-fold from the fetal to the adult stages. Next, to more specifically determine whether such dynamic changes of PcG gene expression occurs in cardiomyocytes (CMs), we isolated neonatal (P0) and adult (3 months) CMs and performed qRT-PCR for 18 PcG genes (**Figure S1A-B**). Among all PcG genes, Cbx7 had the highest fold difference (about 27-fold) between adult and neonatal CMs. In addition, distinct expression patterns were observed such as upregulated (Cbx7, Cbx8, Cbx2, Ring1a and Pcgf5), downregulated (Cbx4, Ezh2, Pcgf2 and Ring1b), and unchanged (Scmh1, Pcgf6, Yy1 and Cbx6). Together, these results imply that CBX7 undergoes the highest increase in the PcG subunit in postnatal CMs.

**Figure 1.**
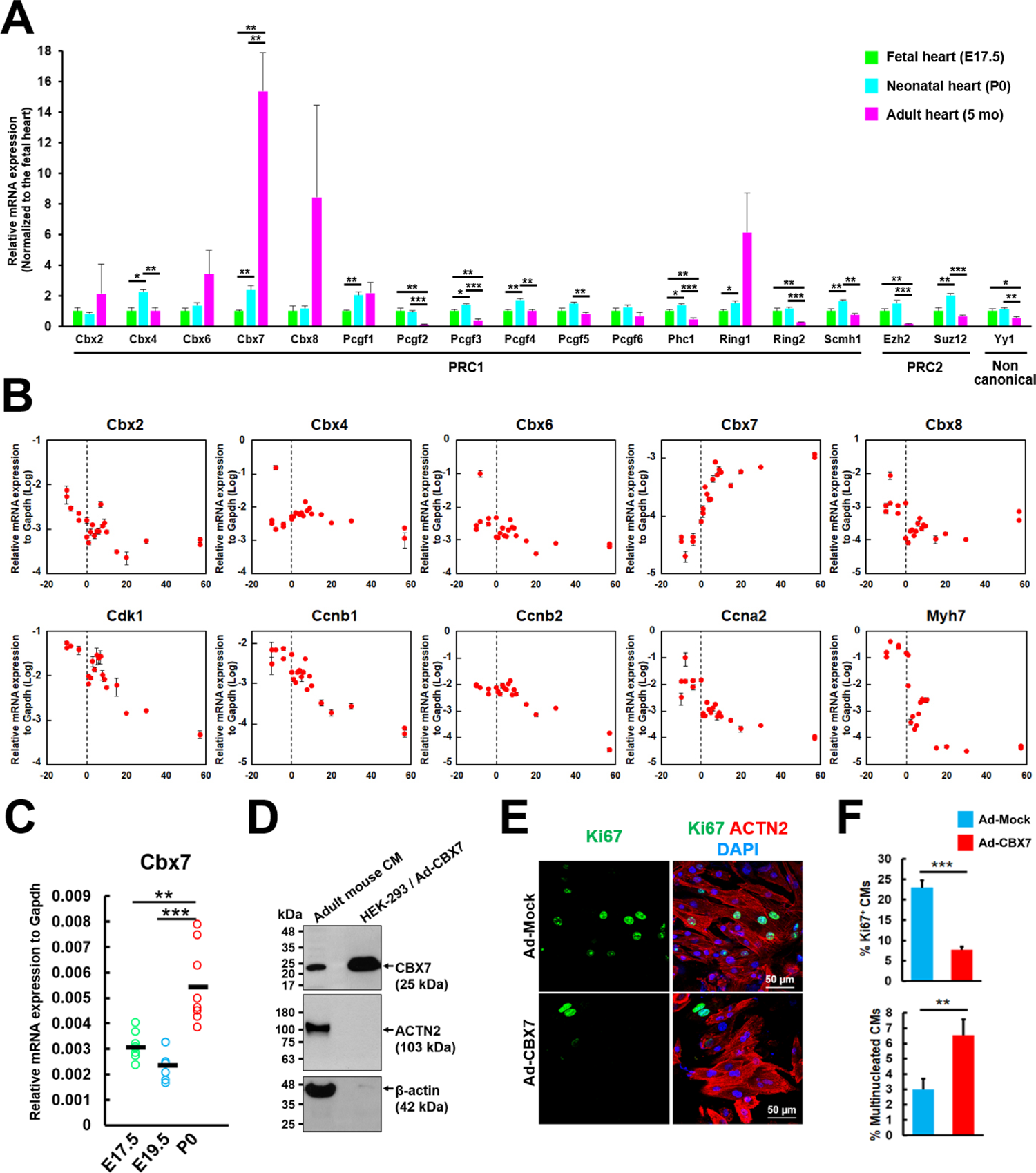
Expression profiling of Polycomb group genes in mouse heart and CMs. **A**. mRNA expression of PcG genes in the mouse heart. Expression of 18 PcG genes were examined in fetal (E17.5), neonatal (P0) and adult (5 months) hearts by qRT-PCR and the fold change was calculated by normalizing to the fetal heart values. *P < 0.05, **P < 0.01, ***P < 0.001. Standard unpaired Student’s t-tests were performed comparing fetal vs. neonatal hearts, neonatal vs. adult hearts, and fetal vs. adult hearts. N = 4, each with technical triplicates. **B**. mRNA levels of the indicated genes in the mouse hearts measured by qRT-PCR at different developmental stages from E10.5 to 60 days after birth. N = 22, each with technical triplicates. **C.** mRNA levels of Cbx7 in mouse hearts showing perinatal upregulation. **P < 0.01, ***P < 0.001. Standard unpaired Student’s t test. N = 8, each with technical triplicates. **D.** Western blot for CBX7 using isolated adult mouse CMs. HEK-293 cells infected with Ad-CBX7 were used as a positive control. **E.** Double immunostaining of neonatal mouse CMs transduced with Ad-CBX7 for ACTN2 and Ki67 and cultured in the presence of growth factors including IGF-1 and FGF-1 for three days. DAPI (blue). **F.** Percentages of Ki67^+^ CMs and multinucleated CMs. *P < 0.05, **P < 0.01, ***P < 0.001. Standard unpaired Student’s t test. More than 500 cells in each group were examined.

### CBX7 expression is increased during the perinatal period and remains high during the postnatal period

Since CBX family genes regulate specific target genes^39^, their expression patterns were further examined in the heart at different time points via qRT-PCR (**Figure 1B**, upper panel). Cbx2 decreased from the fetal to the juvenile period and fluctuated up and down during adulthood. Cbx6 and 8 showed a similar pattern. Cbx4 increased until the preadolescent stage and decreased afterwards. Interestingly, among five mammalian orthologues of CBX proteins (Cbx2, 4, 6, 7 and 8), only Cbx7 showed exponential increase during the perinatal period. Cbx7 expression increased right after birth, peaked at day 6 (at 10-fold higher than the fetal period) and was maintained at a similarly high level during the postnatal period. To examine whether our findings corroborated with previous reports of expression of other genes, we examined expression patterns of genes for cell cycle activators and Myh7, an immature isoform of cardiac myosin heavy chain (**Figure 1B**, lower panel). Consistent with previous studies^40^, multiple Cyclins and CDKs including Ccna2, Ccnb1, Ccnb2, and Cdk1 were downregulated in the postnatal mouse hearts. An immature isoform of cardiac myosin heavy chain Myh7 was also downregulated as the heart matured. These results confirm the quality of our heart samples and consistency with previous observations. To define when Cbx7 expression is induced, fetal (E17.5 and E19.5) and neonatal mouse hearts were subjected to qRT-PCR for Cbx7. mRNA expression of Cbx7 was relatively low during the fetal stage until E19.5 (**Figure 1C**). However, its expression doubled right after birth (postnatal day 0). To determine protein expression of CBX7 in adult mouse (3-month-old) CMs, we performed western blot (**Figure 1D**). The results showed that CBX7 protein is expressed with a molecular weight of 22∼25 kDa, suggesting that p22^CBX7^ is the major isoform present in the postmitotic heart^38^. Human embryonic kidney (HEK)-293 cell lines infected with adenoviral particles inducing overexpression of CBX7 (Ad-CBX7) was used as a positive control (**Figure S2A)**. These data showed that CBX7 transcript and protein are expressed in postnatal CMs.

### CBX7 inhibits proliferation of CMs and cardiac fibroblasts and promotes binucleation of CMs

Next, we determined whether CBX7 overexpression is able to induce proliferation of neonatal mouse CMs. For these gain-of-function analyses, we generated Ad-CBX7 and validated its expression in both HEK-293 cells and mouse embryonic fibroblasts (MEFs) by western blotting (**Figure S2A**). We further confirmed CBX7 expression in neonatal CMs via qRT-PCR (254-fold increase compared to Ad-Mock control) (**Figure S2B**), western blotting (**Figure S2C**) and immunocytochemistry (ICC) (**Figure S2D**). To determine proliferation of neonatal CMs, we added Ad-CBX7 to the cell culture in the presence of growth factors such as IGF-1 and FGF-1 and conducted ICC for Ki67 (**Figure 1E** and **Figure S2E**). CBX7 overexpression resulted in a ∼15% decrease in Ki67^+^ CMs compared to the Ad-Mock group (**Figure 1E**, upper panel); however, the percentage of binucleated CMs was increased in these CMs (**Figure 1E**, lower panel).

The role of CBX7 was considered context- and tissue-selective^36^. Thus, we wondered whether CBX7 represses proliferation of other cardiac cells. To test this idea, we isolated neonatal cardiac fibroblasts, infected them with Ad-CBX7 or Ad-Mock, and performed 3-(4,5-dimethylthiazol-2-yl)-2,5-diphenyltetrazolium bromide (MTT) assay to measure proliferation. CBX7 overexpression resulted in reduced proliferation of fibroblasts up to 60%, suggesting that CBX7 inhibits proliferation of cardiac fibroblasts (**Figure S2F**). Together, these data imply an inhibitory role of CBX7 for proliferation of both CMs and cardiac fibroblasts.

### Generation of mutant mice with targeted inhibition of CBX7 in the cardiomyocyte

To investigate the function of CBX7 in CMs, we used a Cre-loxP recombination system (**Figure S3A**), by which we generated Tnnt2-Cre;Cbx7^floxed/+^ mice by crossing Tnnt2-Cre mice with Cbx7^floxed^ mice (**Figure S3**). In Cbx7^floxed^ mice, exon 2 of the CBX7 gene was replaced by a targeting plasmid encoding a gene trap cassette that consisted of an EN2 splice acceptor, a LacZ reporter gene, internal ribosomal entry site (IRES), a neomycin selection marker, and a SV40 polyA signal (**Figure S3A**). The replacement was mediated by homologous recombination, since the 5’ and 3’ arms of the targeting plasmid were complementary to intergenic regions flanking exon 2 in the Cbx7 allele. The targeted allele was confirmed by conventional PCR using various primers designed to detect specific regions (**Figure S3B**). Through this method, we were able to successfully genotype all the different mutant mouse lines. We next quantified and compared CBX7 transcript levels between the wild-type and the mutant (Tnnt2-Cre;Cbx7^floxed/+^) hearts by qRT-PCR (**Figure S3C**). In the CBX7 haplodeficient mice, CBX7 mRNA levels were half of those in the wild-type mice at postnatal day 0 (P0)^41^. These results indicate that the CBX7 gene was successfully targeted in the mutant mouse line.

### Cardiomyocyte-specific inhibition of CBX7 causes lethality, cardiomegaly, thickening of ventricular walls and impeded myocardial compaction in neonatal mice

Approximately 20% of Tnnt2-Cre;Cbx7^floxed/+^ mice exhibited perinatal mortality (P0-P1) (**Figure S4A)**. The rate of mortality gradually decreased over generations, suggesting reduced penetrance. Those progenies with perinatal death showed abnormal behaviors such as lethargy, inactivity, and repeated convulsions immediately after birth (**Figure 2A** and **Supplementary Video 1**). Such mutant pups succumbed to death within a day (**Supplementary Video 2**). Considering a relatively low expression level of CBX7 in the embryonic heart compared to the adult heart and a previous report of viable adult CBX7 knockout mice^33^, perinatal lethality of cardiac CBX7-haplodeficient mice was unexpected. Viable mutant mice (80%) reached adulthood, but showed reduced fertility. When heterozygotes (Tnnt2-Cre;Cbx7^floxed/+^) were crossed with each other, no viable homozygotes (Tnnt2-Cre;Cbx7^floxed/floxed^) were observed.

**Figure 2.**
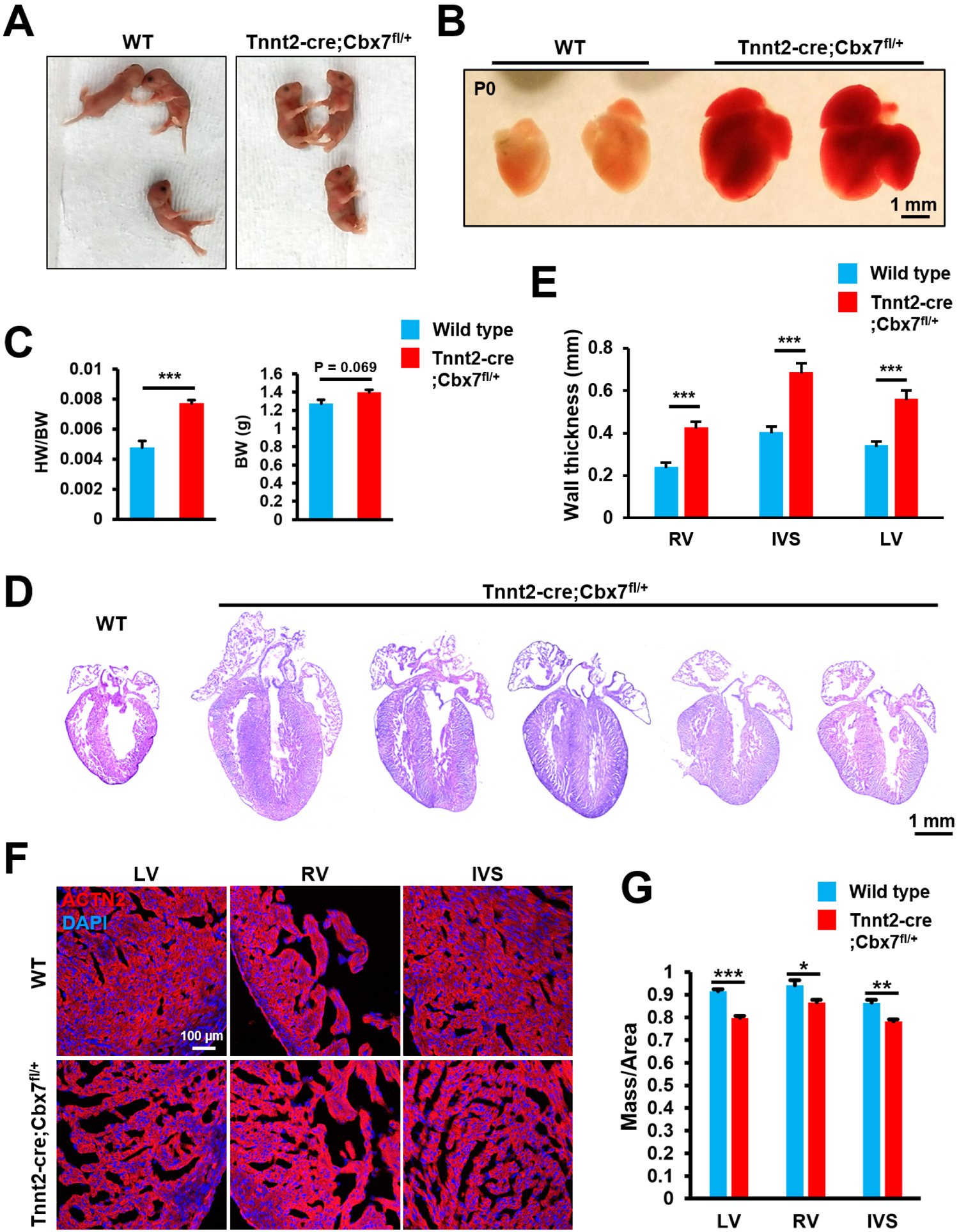
Neonatal lethality, cardiomegaly, thickening of ventricular walls and impeded myocardial compaction by cardiomyocyte-specific inhibition of CBX7. **A.** Representative photographs of neonatal (P0) wild-type and Tnnt2-Cre;Cbx7^fl/+^ mice. **B.** Representative pictures of neonatal (P0) mouse hearts from wild-type and Tnnt2-Cre;Cbx7^fl/+^ mice. **C.** The heart to body weight ratio (left) and the body weight (right). ***P < 0.001. Standard unpaired Student’s t test. N = 3 for the wild-type and N = 10 for the mutant, each with technical triplicates. **D.** H&E stained images of the wild-type and Tnnt2-Cre;Cbx7^fl/+^ hearts in a four chamber view. **E.** Wall thickness of RV, IVS, and LV in wild-type and Tnnt2-Cre;Cbx7^fl/+^. *P < 0.05, **P < 0.01, ***P < 0.001. Standard unpaired Student’s t test. N = 5, each with technical triplicates. **F.** Representative confocal microscopic images of the LV, RV and IVS of control (upper panel) and Tnnt2-Cre;Cbx7^fl/+^ (lower panel) mice at P0 stained for ACTN2 and DAPI. **G.** Quantification of mass per area. *P < 0.05, **P < 0.01, ***P < 0.001. Standard unpaired Student’s t test. N = 3, each with technical triplicates.

To assess any structural defects in cardiac CBX7-haplodeficient mice, hearts were harvested at P0. The mutant mice exhibited cardiomegaly and increased heart weight to body weight ratio (HW/BW) by 38% compared to the WT mice (**Figure 2B-C** and **Figure S4B**) while the body weight was similar between the mutant and WT mice (**Figure 2C**, right). Postmortem samples were excluded because their dehydrated body might affect the results. H & E staining showed substantially increased thickness of ventricular walls (left ventricle (LV), right ventricle (RV), and interventricular septum (IVS)) and enlarged atrium compared to the WT (**Figure 2D and E**). Immunostaining for ACTN2 revealed porous and sponge-like myocardium, a hallmark of fetal heart, and left ventricular non-compaction (LVNC) cardiomyopathy (**Figure 2F**). The mutant showed 8-12% reduced mass per area compared to the WT (**Figure 2G**). Based upon this phenotype at the neonatal stage, we investigated whether CBX7 plays a critical role in cardiac development during the late fetal stage. It is known that cardiac trabeculation occurs at E9.5-E14.5 and myocardial compaction begins at E14.5 and continues through the postnatal stage^42^. We found that the CBX7 transcript was detected in the developing heart at E12.5 and E16.5 (**Figure S4C**). In addition, CM-specific inhibition of CBX7 at E17.5 resulted in more porous and sponge-like myocardium than the control (**Figure S4D**). Together these data indicate that CBX7 represses CM proliferation from the late fetal stage.

### Cardiomyocyte-specific inhibition of CBX7 increased proliferation of neonatal CMs, reduced cell size of neonatal CMs, and induced upregulation of mitotic signaling and downregulation of cardiac maturation-related genes

Since cardiomegaly and ventricular wall thickening seen in the CBX7-haplodeficient mice can be associated with increased proliferation of CMs^43^, we assessed CM proliferation using the neonatal (P0) heart. Double immunostaining for ACTN2 and a proliferation marker, either Ki67 or phospho-histone H3 (pH3), showed expression of Ki67 and pH3 in CMs (**Figure S5A-D**). Quantitatively, double positive cells for ACTN2 and either Ki67 or pH3 were higher in the mutant mice compared to the WT mice; there was a higher percentage of Ki67^+^ CMs than pH3^+^ CMs (**Figure 3A-B**). This is not surprising because pH3 is specific to mitotic cells while Ki67 labels all cycling cells except for resting phase^44^. Since proliferative CMs have smaller CM volume^45^, we also examined the size of CMs by wheat germ agglutinin (WGA) and ACTN2 staining. Compared to the WT, the Cbx7 haplodeficient mice had 2.3-fold smaller CMs (**Figure S6A-B**). These results further support enhancement of CM proliferation by targeted inhibition of CBX7.

**Figure 3.**
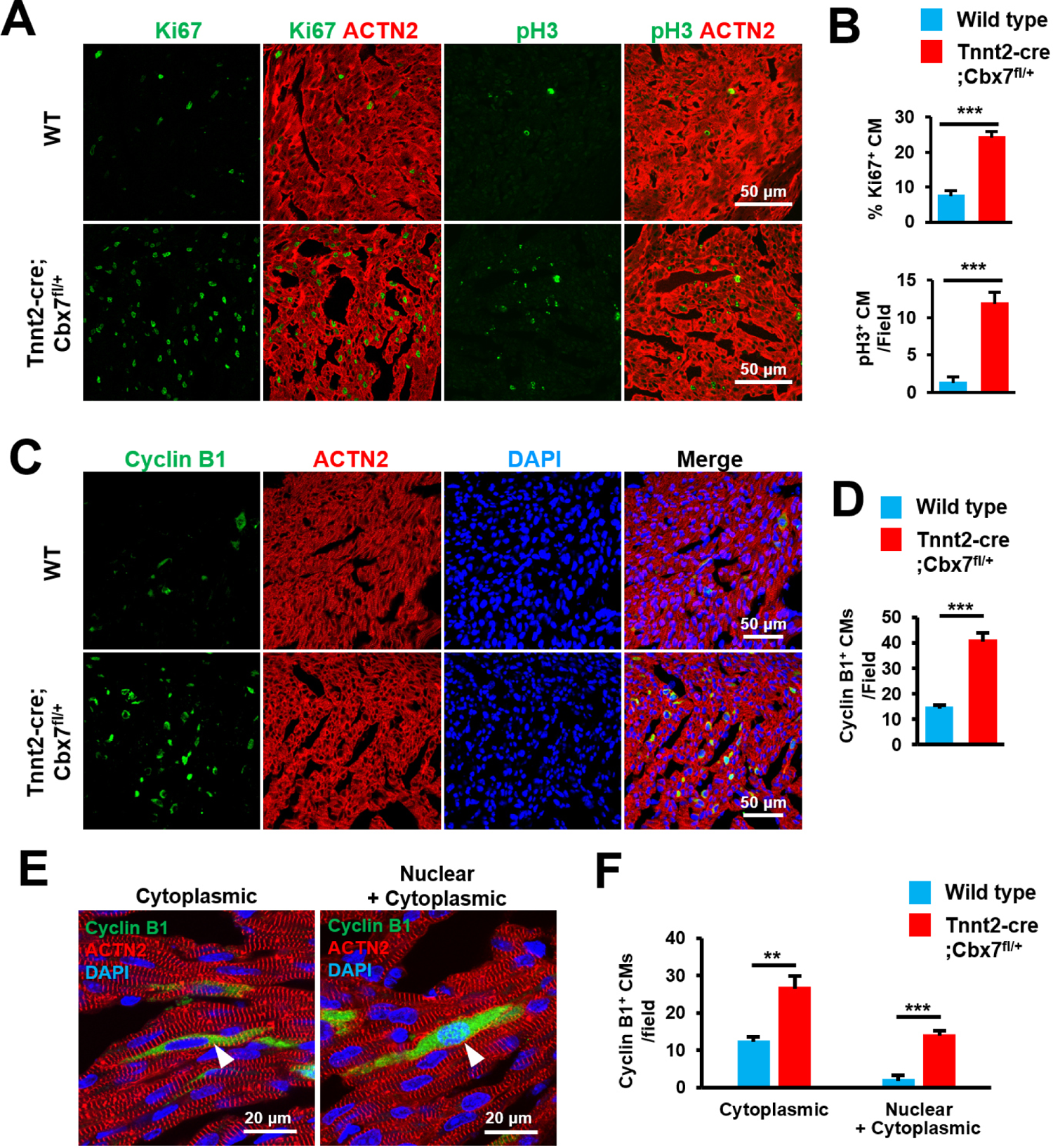
Increased CM proliferation in the neonatal heart by targeted inhibition of CBX7. **A.** Representative confocal microscopic images of neonatal (P0) hearts from wild-type and Tnnt2-Cre;Cbx7^fl/+^ mice stained for ACTN2, Ki67, and pH3. **B.** Quantification of Ki67^+^ or pH3^+^ CMs out of total CMs. ***P < 0.001. Standard unpaired Student’s t test. N = 3. More than 5000 cells in each group were examined. **C-F.** Results from immunostaining of neonatal (P0) hearts from wild-type and Tnnt2-Cre;Cbx7^fl/+^ mice with for Cyclin B1 and ACTN2. DAPI (blue). Representative confocal microscopic images stained for Cyclin B1 and ACTN2 in a low magnification (**C**) and their quantification (**D).** ***P < 0.001, Standard unpaired Student’s t test. N = 3. More than 5000 cells in each group were examined. Representative confocal microscopic images showing two different subcellular localizations of Cyclin B1 in CMs (**E**) and their quantification (**F**). **P < 0.01, ***P < 0.001. Standard unpaired Student’s t test. N = 3. 170 Cyclin B^+^ CMs in 5 different fields were examined.

To further examine the function of CBX7 in mitotic entry of CMs, control and mutant neonatal hearts were immunostained for Cyclin B1. Cyclin B1 functions as a “molecular switch” for mitotic entry. Cyclin B1 is accumulated in the cytoplasm during the whole cell cycle^46, 47^ and upon its phosphorylation, it enters the nucleus and binds to CDK1 to form a functional complex, called “maturation-promoting factor (MPF)”^48, 49^. This complex controls multiple mitotic events such as chromosome condensation, disintegration of nuclear membrane, and assembly of spindle pole^50, 51^. The mutant showed a 2.8-fold increase in the percentage of Cyclin B1^+^ CMs compared to the WT (**Figure 3C-D and Figure S6C**). We further assessed subcellular localization of Cyclin B1 in the WT and mutant CMs and found two different staining patterns of Cyclin B1 in its localization: cytoplasm only and both nuclear and cytoplasm (**Figure 3E**). In both localization patterns, Cyclin B1^+^ CM populations were increased in the CBX7 mutant mice (**Figure 3F**). Together, these data indicate that CBX7 is a critical repressor of CM proliferation at the perinatal stage.

We next investigated changes in the expression of genes associated cell cycle activation and cardiac maturation via qRT-PCR (**Figure S7A-D**). In the mutant hearts, cell cycle activator genes involved in the G2/M phase, Ccna2, Ccnb1, Ccnb2, and Cdk1 were upregulated. However, genes associated with cardiac cytoskeletal/gap junction genes (Myl7, Myl2, Tnnt2, and Gja1) and cardiac ion transporting genes (Kcnj2, Scn5a, and Atp2a2) were downregulated. As for cardiac myofibril maturation, an immature isoform myosin heavy chain Myh7 was upregulated in the mutant hearts but a mature isoform of myosin heavy chain Myh6 was downregulated. As the heart matures, cell cycle activator genes are suppressed in CMs, diminishing proliferative capacity of CMs^40^. On the other hand, genes involved in cardiac maturation are upregulated to meet the metabolic demands of fully functional CMs^52^. Together, these results indicate that genes associated with mitotic signaling were upregulated and those related to cardiac maturation were downregulated by targeted inhibition of cardiac CBX7.

Taken together, genetic inhibition of CBX7 in CMs resulted in increased CM proliferation, reduced CM size, upregulation of cell cycle activator genes, and downregulation of cardiac maturation genes in the mutant heart.

### CBX7 represses proliferation of postnatal CMs

We next examined the role of CBX7 in postnatal CMs. We generated inducible conditional knockout (iCKO) mice targeting CBX7 in CMs (Myh6-MCM;Cbx7^fl/fl^). To avoid potential side effects, we first removed the gene trapping cassette from Cbx7^floxed^ mice by crossbreeding with R26-FLP1 mice (**Figure S8A**). The genotype was confirmed via traditional PCR (**Figure S8B-C**). We designed CBX7 iCKO mice (Myh6-MCM;Cbx7^fl/fl^) to delete exon 2 of the Cbx7 gene upon administration of tamoxifen (**Figure 4A**). We subcutaneously injected tamoxifen to the neonatal mice at P0∼P2 and harvested the hearts at P7 and 3 months (mo) (**Figure 4B**). To validate the depletion of CBX7 protein in the mutant mice, we performed western blotting with 3 mo-hearts using an anti-CBX7 antibody (**Figure 4C**). The CBX7 protein was not completely knocked out but significantly decreased. To measure CM proliferation, P7 hearts were sectioned and immunostained for Ki67, pH3, and Cyclin B1. All these markers were significantly increased in CMs (**Figure 4D-J and Figure S9A**). At 3 months, HW/BW ratio was increased in a subset (20%) of tamoxifen-treated iCKO mice (higher than the mean plus two standard deviations (SD) of the control) which exhibited overt cardiomegaly compared to the vehicle-treated mice (**Figure 4K**), and another subset (40%) exhibited reduced HW/BW ratio (lower than the mean minus two SD of the control) (**Figure S9B**). Ejection fraction measured by echocardiography ranged from 40% to 70% (**Figure S9C**). However, there was no correlation between HW/BW and ejection fraction (**Figure S9D**). There are several possible explanations for this observation. Genetic inhibition of CBX7 in CMs resulted in impeded myocardial compaction (**Figure 2F-G**), which might have reduced cardiac workload, resulting in cardiac atrophy^53^. Another possible cause of cardiac atrophy could be cardiotoxicity of tamoxifen^54^. Finally, CM size was similar between vehicle- and tamoxifen-treated groups (**Figure S8E-F**), suggesting that CBX7 does not affect CM size at adulthood. Together, these data indicated that genetic inhibition of CBX7 in postnatal CMs via tamoxifen-inducible system resulted in increased CM proliferation.

**Figure 4.**
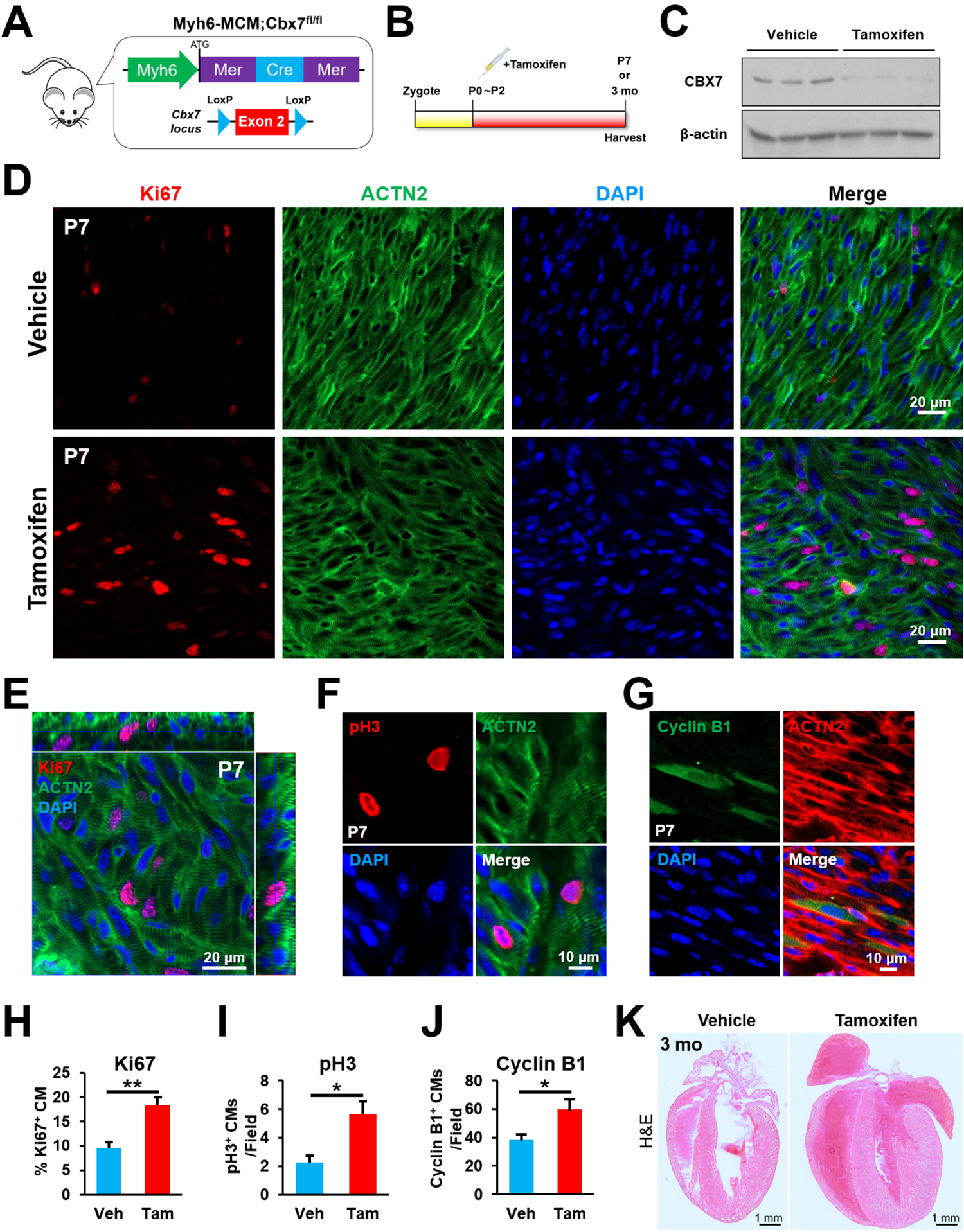
Increased CM proliferation in the postnatal heart by targeted inhibition of CBX7. **A.** A schematic showing the genotype of inducible conditional knockout mice (Myh6-MCM;Cbx7^fl/fl^). Exon 2 of Cbx7 gene is genetically deleted in CMs upon tamoxifen treatment. **B.** The experimental timeline for postnatal deletion of CBX7 in CMs. Tamoxifen was subcutaneously administered to neonatal mice at P0-P2 and the hearts were harvested at P7 or 3 months (mo). **C.** Validation of CBX7 deletion via western blotting with hearts collected from Myh6-MCM;Cbx7^fl/fl^ mice at 3 mo. N = 3, each with technical triplicates. **D.** Representative confocal microscopic images of P7 hearts from Myh6-MCM;Cbx7^fl/fl^ mice immunostained for Ki67, ACTN2. DAPI (blue). **E.** A representative orthogonal projection image of tamoxifen-treated P7 hearts from Myh6-MCM;Cbx7^fl/fl^ mice in panel D. **F.** Representative confocal microscopic images of P7 hearts from Myh6-MCM;Cbx7^fl/fl^ mice immunostained for pH3, ACTN2. DAPI (blue). **G.** Representative confocal microscopic images of P7 hearts from Myh6-MCM;Cbx7^fl/fl^ mice immunostained for Cyclin B1, ACTN2. DAPI (blue). **H-J.** Quantification of CMs positive for Ki67 (H), pH3 (I), and Cyclin B1 (J) in P7 hearts from Myh6-MCM;Cbx7^fl/fl^ mice. *P < 0.05, **P < 0.01. Standard unpaired Student’s t test. N = 3. More than 5000 cells in each group were examined. **K.** Representative H&E stained images of 3 month-old hearts from Myh6-MCM;Cbx7^fl/fl^ mice.

### CBX7 interacts with TARDBP and controls target genes of TARDBP

Previous studies reported that CBX7 recognizes tri-methylated histones and mediates transcriptional repression by interacting with other PcG proteins in the nucleus^37^. Our recent study newly demonstrated that the short isoform of CBX7 (22-25 kDa) exists mainly in the cytoplasm whereas the long isoform (36 kDa) is present in the nucleus^38^. As shown in Figure 1D, the major isoform in adult mouse CMs is the short isoform. When overexpressed in neonatal mouse CMs, it was localized to the cytoplasm and reduced the number of Ki67^+^ CMs (**Figure 1F** and **Figure S2D**). However, the underlying mechanism whereby the short isoform of CBX7 represses cell proliferation in the cytoplasm is unknown. To explore this mechanism, we investigated potential cytoplasmic binding partners of CBX7. Mouse CBX7 (mCBX7) gene was overexpressed in MEFs via adenoviral particles (Ad-mCBX7). The cytoplasmic protein fraction was isolated and subjected to immunoprecipitation using either IgG (control) or anti-CBX7 antibody. Immunoprecipitated samples were subjected to SDS-PAGE followed by silver staining. Two bands were identified, which represent candidates for CBX7 binding partners (**Figure 5A**). Slices for each band underwent matrix-assisted laser desorption/ionization-time of flight (MALDI-TOF) followed by mass spectrometry (MS). Proteins identified with more than 6 exclusive spectrum counts were listed in **Figure 5B**. Among them, we suspected that TAR DNA-binding protein 43 (TARDBP) is a highly likely binding partner for CBX7 since TARDBP showed the highest spectrum counts and its function is related to cell proliferation^55–58^.

**Figure 5.**
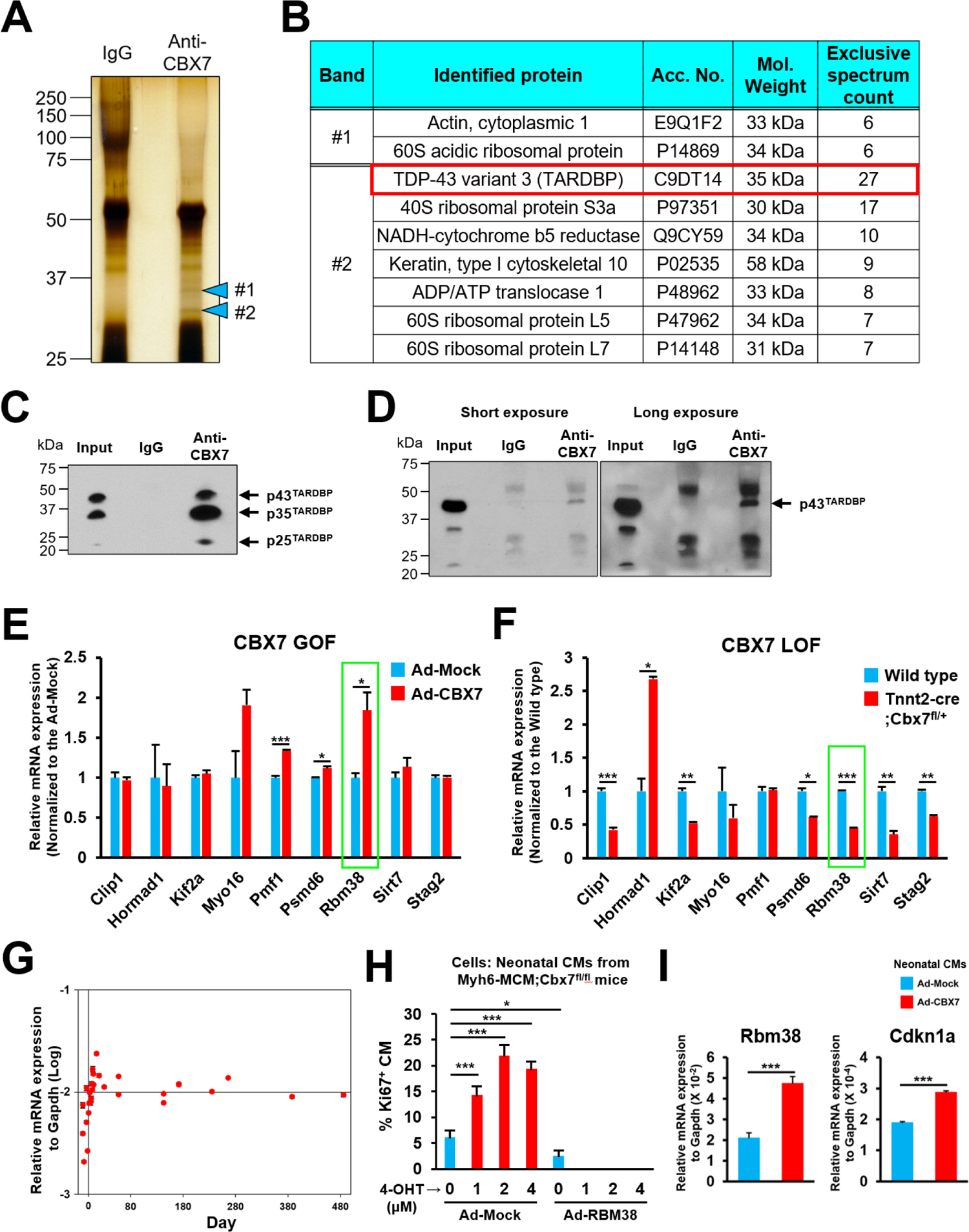
Interaction of CBX7 and TARDBP and its impact on expression of mitosis-related genes. **A**. Immunoprecipitation of CBX7 binding partners. Mouse CBX7 (mCBX7) gene was overexpressed in mouse embryonic fibroblasts (MEFs) via Ad-mCBX7. The cytoplasmic protein fraction was isolated and immunoprecipitated using either IgG (control) or anti-CBX7 antibody. Immunoprecipitated samples were subjected to SDS-PAGE followed by silver staining. Indicated bands (#1 and #2) represent candidates for CBX7 binding partners. **B**. Identified peptides for CBX7 binding partners. Band slices for #1 and #2 underwent MALDI-TOF followed by mass spectrometry. Identified proteins with more than 6 exclusive spectrum counts were listed. Probability for all the listed proteins is higher than 95%. **C-D**. Immunoprecipitation for TARDBP. The cytoplasmic protein fraction was isolated from MEFs treated with Ad-mCBX7 (C) or 3-month-old adult mouse hearts (D) and was immunoprecipitated using either IgG (control) or anti-CBX7 antibody. Immunoprecipitated samples were subjected to SDS-PAGE followed by western blotting with an anti-TARDBP antibody. Input was used as a positive control. N = 3, each with technical triplicates. **E**. mRNA expression of TARDBP target genes in CBX7-overexpressed MEFs. MEFs were treated with Ad-Mock or Ad-CBX7 for three days and underwent qRT-PCR. Gene expression level was normalized to Gapdh. *P < 0.05, ***P < 0.001. Standard unpaired Student’s t test. N = 3, each with technical triplicates. GOF: gain of function. **F**. mRNA expression of TARDBP target genes in CBX7-haplodeficient (Tnnt2-Cre;Cbx7^fl/+^) mice hearts in comparison with neonatal (P0) hearts. Gene expression measured by qRT-PCR and normalized to Gapdh. *P < 0.05, **P < 0.01, ***P < 0.001. Standard unpaired Student’s t test. N = 3, each with technical triplicates. LOF: loss of function **G**. mRNA expression of Rbm38 gene in the mouse hearts from E10.5 to 486 days after birth. qRT-PCR results. N = 31, each with technical triplicates. **H**. Percentages of Ki67^+^ CMs upon CBX7 knockout and Rbm38 overexpression. CMs were isolated from neonatal (P1) Myh6-MCM;Cbx7^fl/fl^ mice and treated with Ad-Mock or Ad-CBX7 in the presence of 4-OHT at different concentrations. Cells were cultured for 5 days in the absence of growth factors such as IGF-1 and FGF-1. *P < 0.05, ***P < 0.001. Standard unpaired Student’s t test. N = 3. More than 500 cells in each group were examined. **I**. mRNA expression of Rbm38 and Cdkn1a genes in neonatal CMs treated with Ad-Mock or Ad-CBX7. Neonatal (P0) mouse CMs were treated with adenoviral particles for 3 days and subjected to qRT-PCR. ***P < 0.001. Standard unpaired Student’s t test. N = 3.

Studies showed that TARDBP is an RNA-binding protein and regulates mitotic signaling by directly binding to mRNAs of cell cycle-related genes^55–58^. It is involved in many RNA processing pathways such as mRNA stability, pre-mRNA splicing, mRNA transport, microRNA processing, transcriptional regulation, translational regulation and stress granule formation^59^. TARDBP interacts with multiple proteins such as 14-3-3, copper/zinc superoxide dismutase (SOD1) and ribosomal proteins^60, 61^, but the interaction with CBX7 has been unknown. To validate the mass spectrometry result, Ad-mCBX7 was transduced into MEFs, and the cytoplasmic protein fraction was isolated and immunoprecipitated with either IgG or anti-CBX7 antibody followed by immunoblotting with an anti-TARDBP antibody. There were three bands at 43 kDa (p43^TARDBP^), 35 kDa (p35^TARDBP^), and 25 kDa (p25^TARDBP^) in the input, indicating three transcript variants^62^(**Figure 5C**). Studies reported that all three isoforms of TARDBP can be localized to both the cytoplasm and the nucleus^62, 63^. When precipitated with anti-CBX7 antibody, all three bands were present, indicating that CBX7 binds to all the three transcript variants of TARDBP in MEFs. To determine this interaction in the postnatal heart, we conducted similar experiments with the cytoplasmic protein fraction from adult mouse hearts (3-months-old). In the input, all the three isoforms of TARDBP were detected but the expression of p43^TARDBP^ was the highest (**Figure 5D**). The control IgG sample showed nonspecific bands at 50 kDa, 30 kDa, and 25 kDa. Immuno-precipitation with an anti-CBX7 antibody showed an additional band at 43 kDa, suggesting that CBX7 interacts with p43^TARDBP^ in the postnatal heart. To further validate this interaction, we performed immunostaining of the adult mouse heart with anti-CBX7 and anti-TARDBP antibodies. CBX7 was colocalized with TARDBP, forming cytoplasmic particles in the postnatal hearts but not in the fetal hearts (**Figure S10**). Similar results were observed in HEK-293 cells treated with sodium arsenite, which is known to induce cytoplasmic TARDBP aggregation (**Figure S11**). These results are consistent with the previous report that p43^TARDBP^ forms particles in the cytoplasm^63^. Together, these results showed that CBX7 can bind to all the alternative splicing isoforms of TARDBP and its major binding partner in the postnatal heart is p43^TARDBP^ in the cytoplasm.

Given the function of TARDBP in mitosis^55–58^, we next investigated whether CBX7 regulates downstream target genes of TARDBP which are associated with mitosis such as CAP-Gly domain containing linker protein 1 (Clip1), HORMA domain containing 1 (Hormad1), kinesin family member 2A (Kif2a), myosin XVI (Myo16), polyamine modulated factor 1 (Pmf1), proteasome 26S subunit non-ATPase 6 (Psmd6), RNA binding motif protein 38 (Rbm38), sirtuin 7 (Sirt7), and stromal antigen 2 (Stag2)^58^ (**Figure 5E-F**). First, we conducted CBX7 gain-of-function (GOF) studies by treating MEFs with Ad-CBX7 particles. qRT-PCR showed that multiple genes were upregulated such as Pmf1, Psmd6, and Rbm38 (**Figure 5E**). Next, we performed CBX7 loss-of-function (LOF) studies by using hearts from neonatal (P0) WT or CBX7 haplodeficient (Tnnt2-Cre;Cbx7^fl/+^) mice. qRT-PCR demonstrated downregulation of all measured genes except Hormad1, which was upregulated (**Figure 5F**). Among the TARDBP target genes we examined, only two genes, Rbm38 and Psmd6, were upregulated by CBX7 GOF and downregulated by CBX7 LOF, suggesting that they are positively regulated by CBX7. Studies showed that Rbm38 represses cell proliferation and its deficiency causes tumorigenesis^64^. Mechanistically, RBM38 binds to the mRNA of CDKN1A and maintains the stability of CDKN1A transcripts, inducing the cell cycle arrest at G1 phase^65^. Psmd6 is a subunit of proteasome 26S which degrades ubiquitinated proteins, regulating a variety of cellular processes such as cell cycle progression and DNA damage repair^66^. Since the function of Rbm38 is more relevant to cell proliferation, we examined its expression pattern in the mouse heart at different time points via qRT-PCR. The results showed that Rbm38 expression increased after birth and was maintained at a similarly high level during the postnatal period (**Figure 5G**). This pattern was similar to that of CBX7 (**Figure 1B**). To determine the role of TARDBP and CBX7 in the regulation of Rbm38, we questioned whether knockdown of TARDBP would reduce Rbm38 expression and whether this reduction can be recovered by CBX7. We utilized siRNA transfection for TARDBP knockdown and adenoviral particles for CBX7 overexpression in HL-1 CMs (**Figure S12**). To validate siRNA transfection, we first transfected siRNA negative control (siNeg) conjugated with a fluorescent dye Cy5 and visualized the localization of transfected siRNA. Two days after transfection, Cy5 signal was detected in most HL-1 CMs (**Figure S12A**). To validate whether transfected siRNA is localized to the cytoplasm or cytoplasmic membrane, we performed confocal microscopy. Cy5 signal was detected inside of the cell (**Figure S12B**) suggesting successful delivery of siRNA to HL-1 CMs. To validate TARDBP knockdown, we performed western blotting and found that the protein level of TARDBP was significantly downregulated by siTARDBP compared to the siNeg (**Figure S12C**). To validate CBX7 overexpression, we treated adenoviral particles (Ad-Mock or Ad-CBX7) and performed qRT-PCR. Gene expression level of CBX7 was upregulated along with Rbm38 and Cdkn1a genes (**Figure S12D**). To examine the role of TARDBP and CBX7 in the regulation of Rbm38 gene expression, we treated siRNAs in combination with adenoviral particles. The knockdown of TARDBP reduced Rbm38 expression (far left vs. middle bars) whereas overexpression of CBX7 recovered the reduced expression of Rbm38 (middle vs. far right bars) (**Figure S12E**). Together, these data suggest that both TARDBP and CBX7 positively regulate the gene expression of Rbm38 and that CBX7 can compensate the downregulation of TARDBP for Rbm38 expression.

To further examine the role of Rbm38 in CM proliferation, we generated adenoviral particles inducing overexpression of RBM38 (Ad-Rbm38). Neonatal CMs isolated from Myh6-MCM;Cbx7^fl/fl^ were treated with either Ad-Mock or Ad-Rbm38 in the presence of 4-hydroxytamoxifen (4-OHT) at different concentrations and cultured without growth factors for five days. Immunostaining of the cells for Ki67 and ACTN2 demonstrated that the proportion of Ki67^+^ CMs increased 2-3-fold after treatment with 4-OHT in the Ad-Mock treated group; however, such increase was not observed in the Ad-Rbm38-overexpressed CMs (**Figure 5H**). Finally, overexpression of CBX7 in neonatal mouse CMs promoted gene expression of Rbm38 and Cdkn1a (**Figure 5I**). These results indicate that Rbm38 is a critical downstream regulator of CBX7 in controlling CM proliferation. Taken together, these data suggest that CBX7 represses CM proliferation via modulation of the TARDBP/Rbm38 pathway (**Figure S12**). This novel mechanism may explain how CM proliferation is regulated during heart development.

## Discussion

Despite extensive efforts^67^, molecular mechanisms of how CMs lose proliferative capacity at the early postnatal stages are poorly understood. In the current study, we identified a novel mechanism of a PcG protein CBX7 in regulating proliferative capacity of CMs. CBX7 expression was dramatically increased during the perinatal period and sustained in the adult heart. Constitutive deletion of CBX7 in CMs led to CM proliferation, thickening of myocardial wall, cardiomegaly, neonatal lethality. Induced deletion of CBX7 in CMs at the neonatal stage (P1) resulted in enhanced CM proliferation at the preadolescent stage (P7) and cardiomegaly in adulthood (3 mo). Overexpression of CBX7 reduced proliferation of neonatal CMs. Mechanistically, CBX7 interacted with TARDBP and positively regulated gene expression of Rbm38, a target of TARDBP, inhibiting CM proliferation.

The present study is the first to demonstrate that a PcG protein CBX7 is a key molecule involved in the perinatal transition of CMs from the proliferative to non-proliferative state. Some signaling pathways such as Hippo, Erbb2, and Meis1 were shown to be induced or lost in the postnatal heart, regulating CM proliferation. However, their roles in perinatal transition of CM proliferation were not reported. In this study, we identified CBX7 as a unique gene that is perinatally induced right after birth and controls CM proliferation. PcG proteins are important transcriptional regulators of the cell cycle and play a crucial role for dynamically regulating the fate of cardiac progenitors during cardiac development^68^. However, their role in CMs and the mechanism of inhibiting cell division were not reported. We focused on the role of a PcG protein CBX7 as it was the most dramatically upregulated gene during the perinatal period, which also sustained in the adult heart among the PcG genes that we examined (Figure 1). CBX7 is ubiquitously expressed in many organs including the heart^27^; however, its perinatal and postnatal upregulation in the heart was unknown. The study of CBX7 was difficult because there was controversy on the role of CBX7 in cell cycle regulation: one promoting cell proliferation and the other inhibiting cell growth. Our recent study gave a clue for this study because we discovered that CBX7 has two different isoforms in vertebrates as p36 and p25 proteins and each plays a different role according to their context and their localization^38^. We found that the p25 isoform is localized in the cytoplasm and represses cell proliferation. In addition, p25 isoform is induced by serum starvation^38^, which generates high levels of reactive oxygen species (ROS), triggering oxidative stress^38, 69^. This understanding recalled us what happens to the heart during the perinatal period. After birth, the heart is suddenly exposed to the oxygen-rich environment and faces oxidative stress, eliciting DNA damage response and cell cycle arrest in CMs^70^. This stress-induced upregulation of CBX7 provides a ground for understanding its role in inhibiting CM proliferation after birth. The necessity of CBX7 in suppressing the proliferative capacity of CMs was further verified by our findings that overexpression of CBX7 reduced proliferation of neonatal CMs. These data suggest that CBX7 can function as a molecular switch to turn off CM proliferation in response to perinatal stress.

A critical role of CBX7 in CM proliferation was clearly demonstrated by the phenotype of the mice with CM-specific inhibition of CBX7 such as perinatal mortality and cardiomegaly. Perinatal lethality of cardiac CBX7-haplodeficient mice was unexpected since Forzati et al showed that CBX7 knockout mice were viable and reached adulthood^33^. This apparent discrepancy could be due to the different knockout strategies between the previous report and our data. Forzati et al deleted exon 5, 6, and 3’UTR whereas we deleted exon 2 that constitutes the chromodomain. It is also possible that infanticide in rodents makes it difficult to observe perinatal lethality in their study. Cardiomegaly and increased thickness of myocardial walls of cardiac CBX7-haplodeficient mice further support an essential role of CBX7 in the heart development (**Figure 2**). In addition, CBX7-haplodeficient myocardium showed sponge-like structure which is a hallmark of the fetal heart in which CMs are proliferative. In other words, the mutant neonatal heart resembled the normal fetal heart although the heart size is bigger. This implies that the proliferative state of the fetal heart was prolonged to the neonatal stage when CBX7 is inhibited. The percentage of Ki67^+^ or pH3^+^ CMs was significantly increased in the CBX7-haplodeficient hearts. There were higher percentage of Ki67^+^ CMs than pH3^+^ CMs because pH3 is specific to mitotic cells whereas Ki67 can be expressed in all active cell cycle stages except for the resting phase^71–73^. We also observed that targeted inhibition of CBX7 in CMs resulted in increased number of Cyclin B1^+^ CMs. Mohamed et al showed that Cyclin B1 is one of the most crucial factors for inducing CM division^74^. Studies also reported that the neonatal heart exhibited downregulation of multiple cell cycle-related proteins such as Cyclin A, Cyclin B, Cyclin D1-3, CDK1, CDK2, CDK4, and CDK6, compared to the fetal heart^75–81^. Among them, Cyclin B was repressed at a narrow perinatal window between E19 and P1^82^. Our data showed that CBX7 was induced at this stage (Figure 1) and its targeted inhibition led to derepression of Cyclin B1 (Figure 3C-F), indicating that CBX7 played an important role in the repression of Cyclin B1 in CMs during the perinatal period. Together, these data strongly support that CBX7 can function as a major regulator checking CM proliferation as the heart matures during the late developmental stage.

We further uncovered a mechanism regulating CM proliferation via CBX7-TARDBP axis. TARDBP is widely expressed in various tissues including heart, lung, liver, spleen, kidney, and brain and binds to a wide spectrum of RNAs, regulating mRNA stability, pre-mRNA splicing, mRNA transport, microRNA processing, transcriptional regulation, translational regulation and stress granule formation^59^. In addition, TARDBP controls mitotic cell cycle by interacting with mRNAs of cell cycle-related genes^55–58^. Previously identified binding partners for TARDBP include SOD1 and ribosomal proteins^60, 61^. Here, we for the first time identified CBX7 as an interacting partner for TARDBP. TARDBP exists as three alternative splicing isoforms, p43^TARDBP^, p35^TARDBP^ and p25^TARDBP^^62^. Our study showed that CBX7 can interact with all three isoforms of TARDBP and its major binding partner in the postnatal heart was p43^TARDBP^. We also showed that CBX7 affects gene expression of multiple target mRNAs of TARDBP, including Rbm38. RNA-binding proteins can induce substantial changes in gene expression by inducing conformational changes of mRNA, converting the mRNA into stable or unstable states^83^. Thus, we speculate that binding of CBX7 causes conformational changes of TARDBP and its associated mRNA, leading to increased Rbm38 mRNA. RBM38 is also an RNA-binding protein and is known to repress cell proliferation by binding to the mRNA of CDKN1A and maintaining the stability of CDKN1A transcripts, inducing the cell cycle arrest^65^. Consistently, our data showed that overexpression of RBM38 repressed proliferation of neonatal CMs. In addition, gene expression of Rbm38 and Cdkn1a was upregulated by overexpression of CBX7 in neonatal CMs, indicating that Rbm38 is a critical target of CBX7/TARDBP for CM cell cycle arrest. Our data support our working model where CBX7 inhibits CM proliferation by interacting with TARDBP and thereby upregulating Rbm38/Cdkn1a during the perinatal period.

We identified CBX7 as a novel repressor of CM proliferation functioning during the perinatal period. Our data indicate that postnatal upregulation of CBX7 is responsible for the loss of proliferative capacity of CMs. In addition, we revealed a previously unknown molecular mechanism whereby CBX7 regulates TARDBP/Rbm38 axis. This study provides insights into cardiac development in the context of CM proliferation.

## Acknowledgments

We gratefully acknowledge the Emory Microscopy in Medicine (MiM) Core and the Emory Children’s Pediatric Research Center flow cytometry core. This work was supported by grants from NHLBI (R01HL127759, R01HL150877, R61HL 154116,) and NIDDK (DP3-DK108245), and grants from the Bio & Medical Technology Development Program of the National Research Foundation (NRF) funded by the Korean government (MSIP) (No.2020M3A9I4038454, No. 2020R1A2C3003784). K. C. is a recipient of an American Heart Association predoctoral fellowship grant (16PRE31350034). We gratefully acknowledge Dr. Ho-Wook Jun and Dr. Patrick Tj Hwang for mass spectrometry analyses.

## Author Contributions

Conceptualization, K. C., Y.-S. Y., and S. B.; Investigation, K. C. and M. A.; Writing - Original Draft, K. C.; Writing - Review & Editing, K. C., Y.-S. Y., A. H., R. L. S., J. W. C. and C. P.; Project Administration, S. K., J. K., and E. Y. J.; Funding Acquisition, Y.-S. Y.

## Experimental Procedures

### Mice, Breeding and Genotyping

Mice were used in accordance with animal protocols approved by the Emory University Institutional Animal Care and Use Committee (IACUC). Tg(Tnnt2-Cre)5Blh/JiaoJ, B6.FVB(129)-A1cfTg(Myh6-Cre/Esr1*)1Jmk/J (Myh6-MCM), and 129S4/SvJaeSor-Gt(ROSA)26Sortm1(FLP1)Dym/J (R26-FLP1) mice were purchased from the Jackson Laboratories. ICR CD-1 mice were purchased from the Charles River Laboratories. Cbx7^tm1a(KOMP)Wtsi^ mouse was generated by the trans-NIH Knock-Out Mouse Project (KOMP). Its sperm was obtained from the KOMP Repository and rederived via IVF and transplantation to B6 wildtype donor female mice by the Mouse Transgenic and Gene Targeting Core (TMF) at Emory University. Primers used for genotyping are listed in **Supplementary table 1**. Purchased or rederived mice were crossed with B6 wildtype mice. Heterozygotes for either Tg(Tnnt2-Cre)5Blh/JiaoJ or Cbx7^tm1a(KOMP)Wtsi^ allele were selected from F1 progenies and outcrossed with each other to generate cardiac Cbx7-haplodeficient mice (*Tnnt2-Cre;Cbx7^floxed/+^*). Cbx7^floxed^ mice were crossbred with R26-FLP1 mice to remove the gene trapping cassette containing EN2SA, β-gal, IRES, Neo, and SV40pA. The progeny lacking the gene trapping cassette were crossbred with Myh6-MCM mice. Myh6-MCM;Cbx7^fl/+^ mice were bred to each other to produce Myh6-MCM;Cbx7^fl/fl^. Tamoxifen was administered into the neonatal mice as described previously^84^.

## Histological analysis and immunocytochemistry

After euthanasia, mouse heart tissues were removed, fixed in 2% paraformaldehyde (PFA) at 4°C for 16 hours, and submerged in 30% sucrose solution at 4°C for 24 hours. Frozen heart sections prepared with OCT compound (Tissue-TeK 4583) were washed with PBS, and then permeabilized/blocked with PBS containing 0.5% Triton X-100 and 2.5% BSA at room temperature for 1hr. Slides were then incubated with anti-ACTN2 (Sigma, A7811, 1:100), anti-CBX7 (Abcam, ab21873, 1:100), anti-Ki67 (Cell Marque, 275R-14, 1:100), anti-phospho-Histone H3 (Ser10) (Millipore, 06-570, 1:100), anti-Cyclin B1 (Cell Signaling Technology, 12231, 1:100), anti-Ubiquityl-Histone H2A (Lys119) (Cell Signaling Technology, 8240, 1:100), anti-TDP43 (TARDBP) (R&D systems, 982022, 1:100), or anti-mTOR (Cell Signaling Technology, 2983, 1:100) at 4°C overnight. WGA staining was performed by following the manufacturer’s instructions (Thermo Fisher, W32466). The slides were washed three times with PBS containing 0.1% Tween 20 and incubated with appropriate secondary antibodies at room temperature for 1-2 hours. DAPI was used for nuclear staining. The samples were visualized under a Zeiss LSM 510 Meta confocal laser scanning microscope (Carl Zeiss). H&E-stained samples were examined under an Olympus IX71 bright field microscope and a Zeiss Axioskop 2 Plus. For immunocytochemistry, cells were fixed in 4% PFA at room temperature for 10 minutes. Then, samples were permeabilized/blocked with PBS containing 0.1% Triton X-100 and 2.5% BSA at room temperature for 1hour. The rest of procedure is the same as described above for immunohistochemistry.

## Isolation and culture of cardiac cells

Adult mouse CMs were isolated via the conventional Langendorff method as described previously^85^ and used for ICC, WB, and qRT-PCR. For neonatal CM isolation, we modified a previously reported protocol^86^. Briefly, neonatal hearts from P0 mice were minced with scissors and incubated in HBSS containing 0.0125% trypsin overnight at 4°C for pre-digestion. On the following day, the tissue fragments were further digested with the Neonatal Heart Dissociation Kit (Miltenyi Biotec, 130-098-373). After filtering with a 100 µm strainer, cells were plated and cultured in Plating medium (84% DMEM high glucose, 10% horse serum, 5% FBS, 1% anti-anti) at 37 °C for 1.5 hrs, allowing the preferential attachment of fibroblasts. Adherent cells were sub-cultured and used for cardiac fibroblasts which were cultured in DMEM high glucose media containing 10% FBS. Non-adherent cells were subjected to percoll-based separation^87^ and MACS using feeder removal microbeads (Miltenyi Biotec, 130-095-531) for the further purification of neonatal CMs. Enriched neonatal CMs were plated onto collagen-coated dishes and cultured in Plating medium. On the following day, the medium was changed to the Culture medium (78% DMEM high glucose, 17% M-199, 4% horse serum, 1% anti-anti, 1% ITS).

Growth factors including 50 ng/ml IGF1 (BioLegend, 591402) and 25 ng/ml FGF1 (BioLegend, 750902) were added to both Plating and Culture media. Two days after the treatment with recombinant adenoviral particles, cells were subjected to ICC.

## Cell culture

Mouse embryonic fibroblasts (MEFs) were cultured in DMEM high glucose media containing 10% FBS. HL-1 mouse CM cell line was cultured in Claycomb Medium, supplemented with 100 μM norepinephrin, 10% fetal bovine serum (FBS) and 4 mM L-glutamine. The negative control for siRNA and siTARDBP were purchased from Sigma Aldrich (SIC003 and EMU-219331, respectively). We transfected siRNAs by using Lipofectamine™ RNAiMAX Transfection Reagent (Thermo Fisher, 13778100), following the manufacturer’s instruction.

## Generation of recombinant adenovirus particles

Molecular cloning of Cbx7 cDNA was previously described^38^. Rbm38 cDNA was purchased from Genecopoeia (EX-Mm34611-M83). Cbx7 and Rbm38 cDNAs were subcloned into an adenoviral shuttle plasmid, pDC316 (Microbix Biosystems, Mississauga, ON, Canada). Both adenoviral genomic and shuttle plasmids were transfected into HEK-293 cells using Lipofectamine 3000 (Thermo Fisher Scientific, Waltham, MA, USA). Recombinant adenoviral particles were extracted from cell lysates and the titers of adenoviral particles were determined by counting infected colonies using an antibody-mediated detection method (Clontech, Mountain View, CA, USA, Cat# 632250).

## Quantitative Real-time PCR

Total RNA from mouse CMs and hearts were isolated using a guanidinium extraction method^88^ combined with an RNA extraction kit (Qiagen). Extracted RNA was reverse-transcribed using Taqman Reverse Transcription Reagents (Applied Biosystems, 4304134) according to the manufacturer’s instructions. The synthesized cDNA was subjected to qRT-PCR using specific primers and probes (see **Supplementary Table 1**). Quantitative assessment of RNA levels was performed using an ABI PRISM 7500 Sequence Detection System (Applied Biosystems). Relative mRNA expression was normalized to Gapdh.

## Western blot

Cells were lysed with RIPA buffer, supplemented with PMSF, phosphatase-inhibitor cocktail (Sigma) and protease-inhibitor cocktail on ice for 1 h and the lysates were clarified by centrifugation. Equal amounts of lysates were subjected to SDS-PAGE, transferred onto a polyvinylidene fluoride (PVDF) membrane, and blocked for 1 h at room temperature in Tris-buffered saline with 0.05% Tween-20 (TBST) and 5% non-fat milk. The membrane was subsequently incubated with anti-CBX7 (Abcam, ab21873, 1:1000), anti-β-actin (Cell Signaling Technology, 4967, 1:1000), anti-ACTN2 (Sigma, A7811, 1:1000), and anti-TDP43 (TARDBP) (R&D systems, 982022, 1:2500) at 4°C overnight. After washing with TBST, blots were incubated with the appropriate secondary antibodies for 1 h at room temperature and developed using ECL detection reagent (Thermo Fisher, 32106).

## MTT assay

Neonatal (P0) cardiac fibroblasts were seeded on 96 well plate at 5 x 10^3^ cells per well. DMEM high glucose media supplemented with 10% FBS, 1% glutamine, 1% non-essential amino acid (NEAA) was used for culture. Two hours after seeding, cells were treated with adenoviral particles and further incubated for three days. MTT reagent was added to cell culture at a final concentration of 0.5 mg/ml. The plate was incubated at 37°C for 30 min in the dark. After removal of culture media, cells were lysed by DMSO and color was measured at 570 nm.

## Immunoprecipitation, silver staining, and mass spectrometry

Cytoplasmic protein fractions were extracted from MEFs or 3-month-old adult mouse hearts using NE-PER Nuclear and Cytoplasmic Extraction Reagents (Thermo Fisher, 78835). For immunoprecipitation, protein samples were pre-cleared using Dynabeads Protein G (Thermo Fisher, 10004D, 1.5 mg). Primary antibodies, rabbit IgG (Abcam, ab37415, 3 µg) or anti-CBX7 antibody (Abcam, ab21873, 3 µg), were added to the lysates and incubated at 4°C overnight with gentle agitation. To pull down protein:antibody complexes, 1.5 mg of Dynabeads Protein G was added and incubated at 4°C for 3 hrs with gentle agitation. Bead:protein:antibody complexes were washed with ice-cold PBS 6 times and denatured at 95°C for 10 min in Laemmli sample buffer supplemented with 2.5% β-Mercaptoethanol. Supernatants containing protein and antibody were used for SDS-PAGE. Silver staining was performed using Perce Silver Stain Kit (Thermo Fisher, 24612). Gel slices were reduced, carbidomethylated, dehydrated, and digested with Trypsin Gold (Promega). Following digestion, peptides were extracted and all fractions were combined together. The combined fractions were concentrated using a SpeedVac to near dryness, and then resuspended to 20 µl using 95% ddH2O/5% acetonitrile (ACN)/0.1% formic acid (FA) prior to analysis by 1D reverse phase LC-nESI-MS2.

For Mass Spectrometry (MS), peptide digests (8µL each) were injected into a 1260 Infinity nHPLC stack (Agilent) and separated using a 75 micron I.D. x 15 cm pulled tip C-18 column (Jupiter C-18 300 Å, 5 micron, Phenomenex). This system was operated in-line with a Thermo Orbitrap Velos Pro hybrid mass spectrometer, which was equipped with a nano-electrospray source (Thermo Fisher Scientific). All data were collected in collision-induced dissociation (CID) mode. The nHPLC ran with binary mobile phases that included solvent A (0.1% FA in ddH2O), and solvent B (0.1% FA in 15% ddH2O/85% ACN), programmed as follows; 10 mins at 0% solvent B (2 µL/min, load), 90 mins at 0%-40% solvent B (0.5nL/min, analyze), 15 mins at 0% solvent B (2µL/min, equilibrate). Following each parent ion scan (350-1200 m/z at 60k resolution), fragmentation data (MS2) was collected on the top most intense 15 ions. For data dependent scans, charge state screening and dynamic exclusion were enabled with a repeat count of 2, repeat duration of 30 secs, and exclusion duration of 90 secs.

For MS data conversion and searches, the XCalibur RAW files were collected in profile mode, centroided and converted to MzXML using ReAdW v.3.5.1. The mgf files were then created using MzXML2 Search (included in TPP v.3.5) for all scans. The data was searched using SEQUEST, which was set for two maximum missed cleavages, a precursor mass window of 20 ppm, trypsin digestion, variable modification C at 57.0293, and M at 15.9949. Searches were performed with a species-specific subset of the UniRef100 database.

For further data analyses, the list of peptide IDs generated based on SEQUEST search results were filtered using Scaffold (Protein Sciences, Portland Oregon). Scaffold filters and groups all peptides to generate and retain only high confidence IDs while also generating normalized spectral counts (N-SC’s) across all samples for the purpose of relative quantification. The filter cut-off values were set with minimum peptide length of >5 AA’s, with no MH+1 charge states, with peptide probabilities of >80% C.I., and with the number of peptides per protein ≥2. The protein probabilities were then set to a >99.0% C.I., and an FDR<1.0. Scaffold incorporates the two most common methods for statistical validation of large proteome datasets, the false discovery rate (FDR) and protein probability^89–91^. Relative quantification across experiments were then performed via spectral counting^92, 93^, and when relevant, spectral count abundances were normalized between samples^94^.

## Echocardiography

Echocardiography was performed on adult mice (3-month-old) using the Vevo 3100^TM^ Imaging System (VisualSonics, Inc) as previously described^95^. Ejection fraction (EF) was measured using a two-dimensional method.

## Statistical analyses

Investigators were blinded to the assessment of the analyses of cell, animal, and histological experiments. Both sexes were included in the animal data. For the quantification of Ki67, pH3, and Cyclin B1^+^ CMs and cell size (WGA), the results were obtained from 5 sections of the heart from each animal. All data were presented as mean ± standard error of the mean (SEM). For statistical analysis, the standard unpaired Student’s t-test and one-way ANOVA test with Tukey HSD were performed. For the correlation analysis, Pearson’s correlation coefficient was calculated. P values less than 0.05 were considered statistically significant.

