## Supplementary Data for "Polycomb group protein CBX7 represses cardiomyocyte proliferation via modulation of the TARDBP/Rbm38 axis"

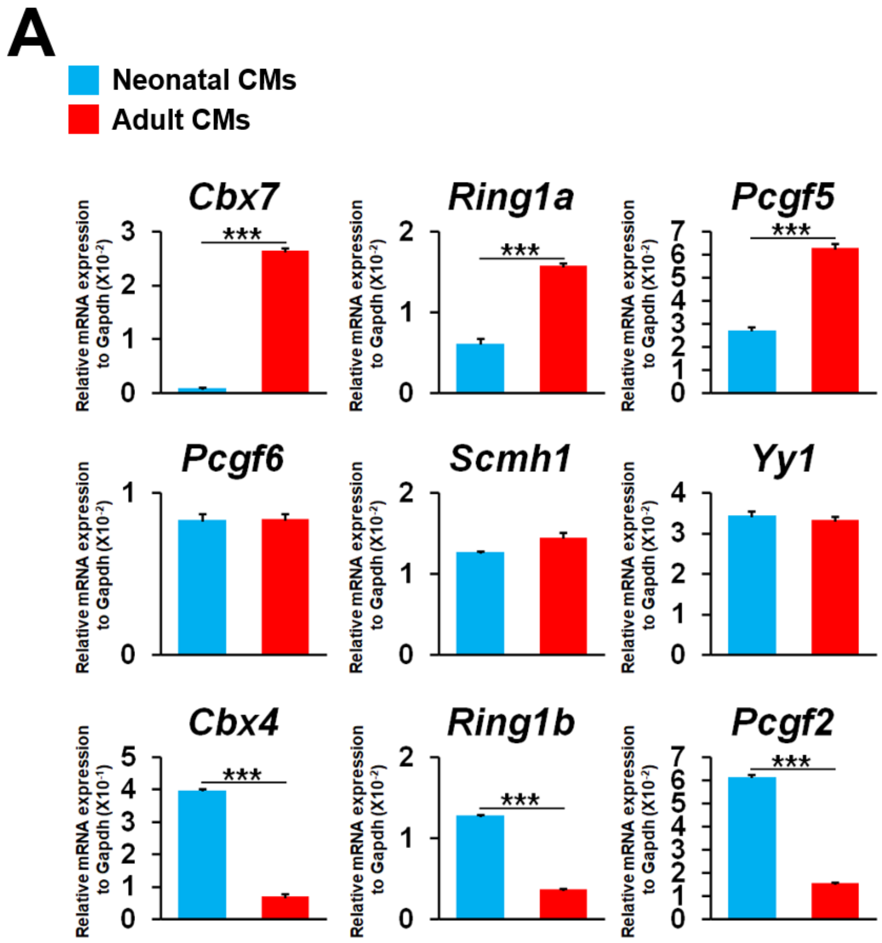


**B**

| Adult / Neonate | |
|---|---|
| **Gene** | **Fold Change** |
| ***Cbx7*** | **27.32** |
| ***Cbx8*** | **6.48** |
| ***Cbx2*** | **2.86** |
| ***Ring1a*** | **2.56** |
| ***Pcgf5*** | **2.32** |
| ***Pcgf1*** | **1.93** |
| ***Pcgf4*** | **1.40** |
| ***Scmh1*** | **1.13** |
| ***Pcgf6*** | **1.00** |
| ***Yy1*** | **-1.04** |
| ***Cbx6*** | **-1.04** |
| ***Phc1*** | **-1.22** |
| ***Pcgf3*** | **-1.81** |
| ***Suz12*** | **-1.97** |
| ***Ring1b*** | **-3.48** |
| ***Pcgf2*** | **-3.93** |
| ***Ezh2*** | **-4.79** |
| ***Cbx4*** | **-5.67** |

**Figure S1. Gene expression profiling of Polycomb group proteins in mouse CMs. A.** Representative qRT-PCR results. Expression patterns of 18 *PcG* genes were examined in primary neonatal (P0) and adult (3 months) mouse CMs by qRT-PCR. ***P < 0.001. Standard unpaired Student's t test. N = 3, each with technical triplicates. **B.** qRT-PCR results of 18 PcG genes expressed as a ratio of adult CMs vs. neonatal CMs.

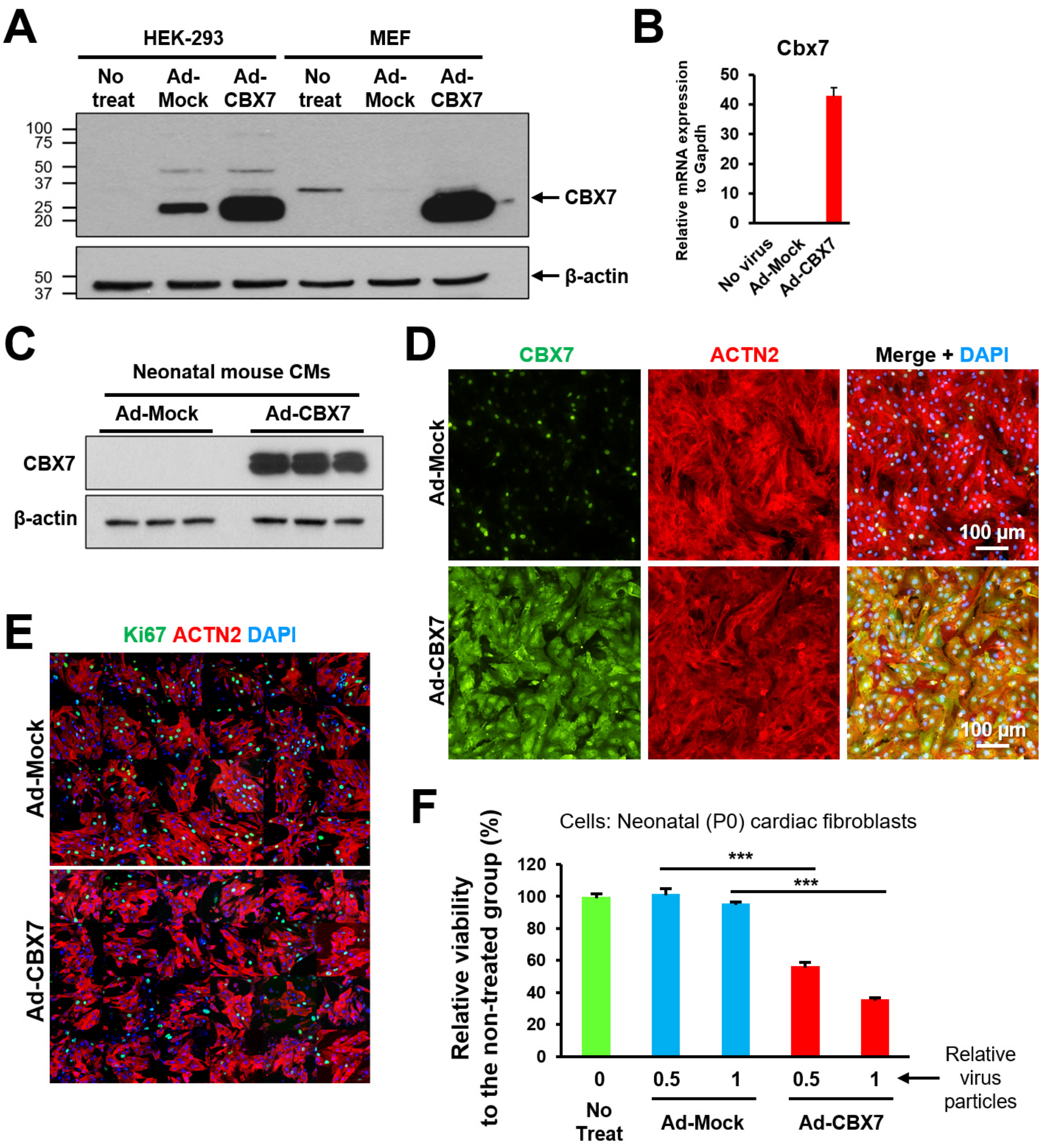


**Figure S2. CBX7 gain-of-function analyses in neonatal mouse CMs and cardiac fibroblasts.** **A.** Validation of CBX7 overexpression via western blotting. HEK-293 and MEFs were treated with nothing (No treat), empty adenoviral particles (Ad-Mock) or adenoviral particles inducing mouse CBX7 overexpression (Ad-CBX7). Three days after transduction, whole cell lysates were subjected to western blot. **B-C**. Validation of CBX7 overexpression in neonatal CMs. Neonatal

(P0) mouse CMs were treated with adenoviral particles for 3 days and subjected to qRT-PCR (**B**) or western blotting (**C**) for CBX7. N = 3, each with technical triplicates. **D.** Representative epifluorescence microscopic images of neonatal (P0) mouse CMs immunostained for CBX7 and ACTN2. Neonatal (P0) mouse CMs were infected with Ad-Mock or Ad-CBX7 adenoviruses for 3 days. DAPI (blue). **E.** Low magnification images used for quantification in Figure 1E. Three days after adenoviral transduction, neonatal mouse CMs were immunostained for ACTN2 and Ki67. DAPI (blue). **F.** MTT assay results for neonatal mouse cardiac fibroblasts infected with either Ad-Mock or Ad-CBX7 viral particles for 3 days with different titers. ***$P < 0.001$. Standard unpaired Student's t test. N =3, each with technical triplicates.

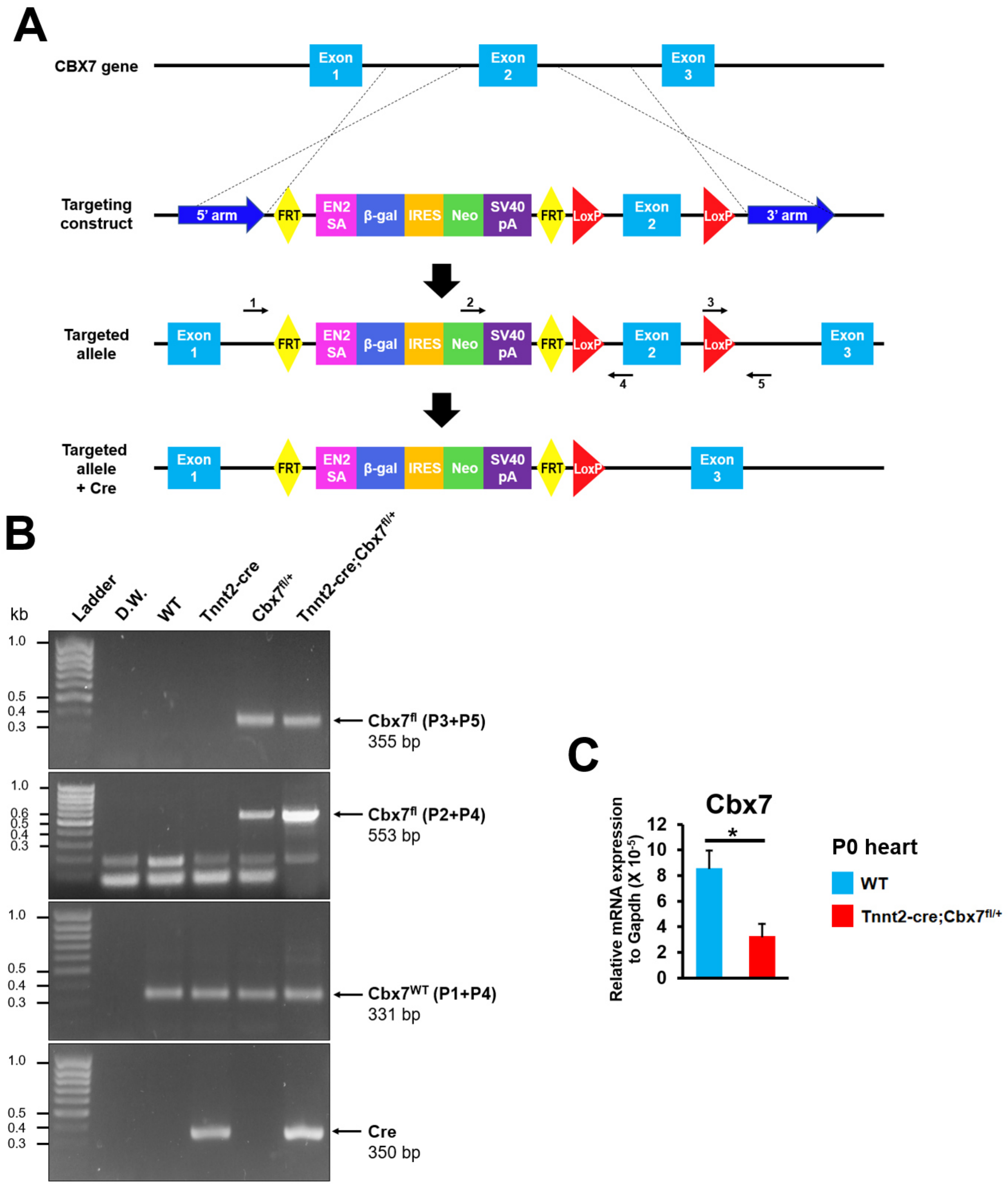


**Figure S3. A schematic and validation of the targeted deletion of a Cbx7 allele. A.** A schematic showing a Cbx7 allele, the targeting construct, and the targeted allele. Arrows with numbers represent primers specifically designed to the indicated regions. **B.** Conventional PCR results showing different target sequences in a CBX7 allele using genomic DNA collected from the indicated mice. **C.** Representative qRT-PCR for Cbx7 using cDNA from wild-type and Tnnt2-Cre;Cbx7$^{fl/+}$ neonatal hearts (P0). *P < 0.05. Standard unpaired Student's t test. N = 3, each with technical triplicates.

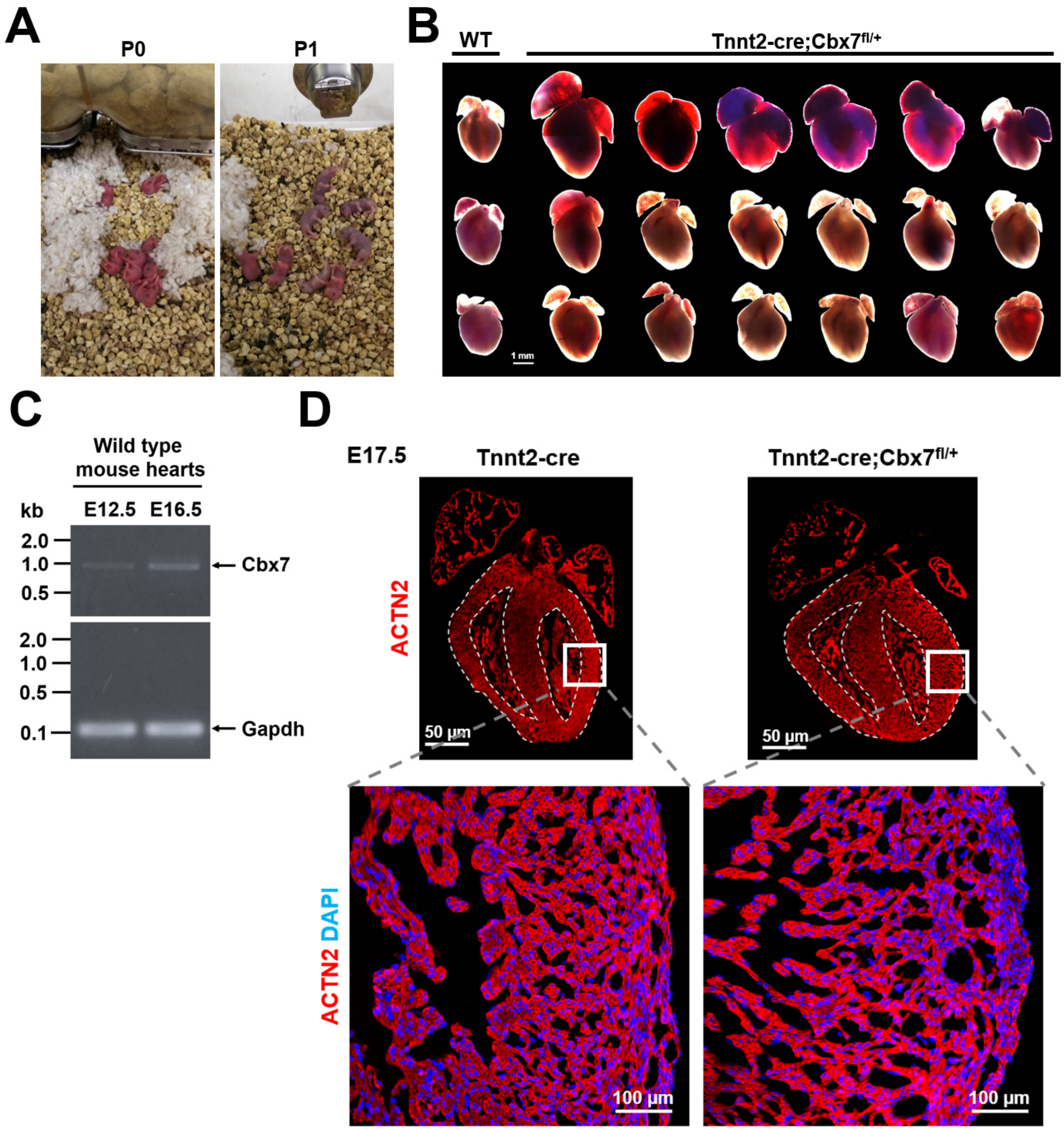


**Figure S4. Neonatal lethality with increased heart size and impeded myocardial compaction in fetal mouse hearts by targeted inhibition of CBX7. A.** Representative photographs of Tnnt2-Cre;Cbx7$^{fl/+}$ mice at P0 (left) and P1 (right). **B.** Representative bright field images of neonatal (P0) hearts from wild-type and Tnnt2-Cre;Cbx7$^{fl/+}$ mice. **C.** RT-PCR results for a CBX7 transcript (full length) with fetal mouse hearts at E12.5 and E16.5. **D.** Representative confocal microscopic images of hearts immunostained for ACTN2 and DAPI. Fetal heart at E17.5 (upper panel) and left ventricle (lower panel) of control and Tnnt2-Cre;Cbx7$^{fl/+}$ mice.

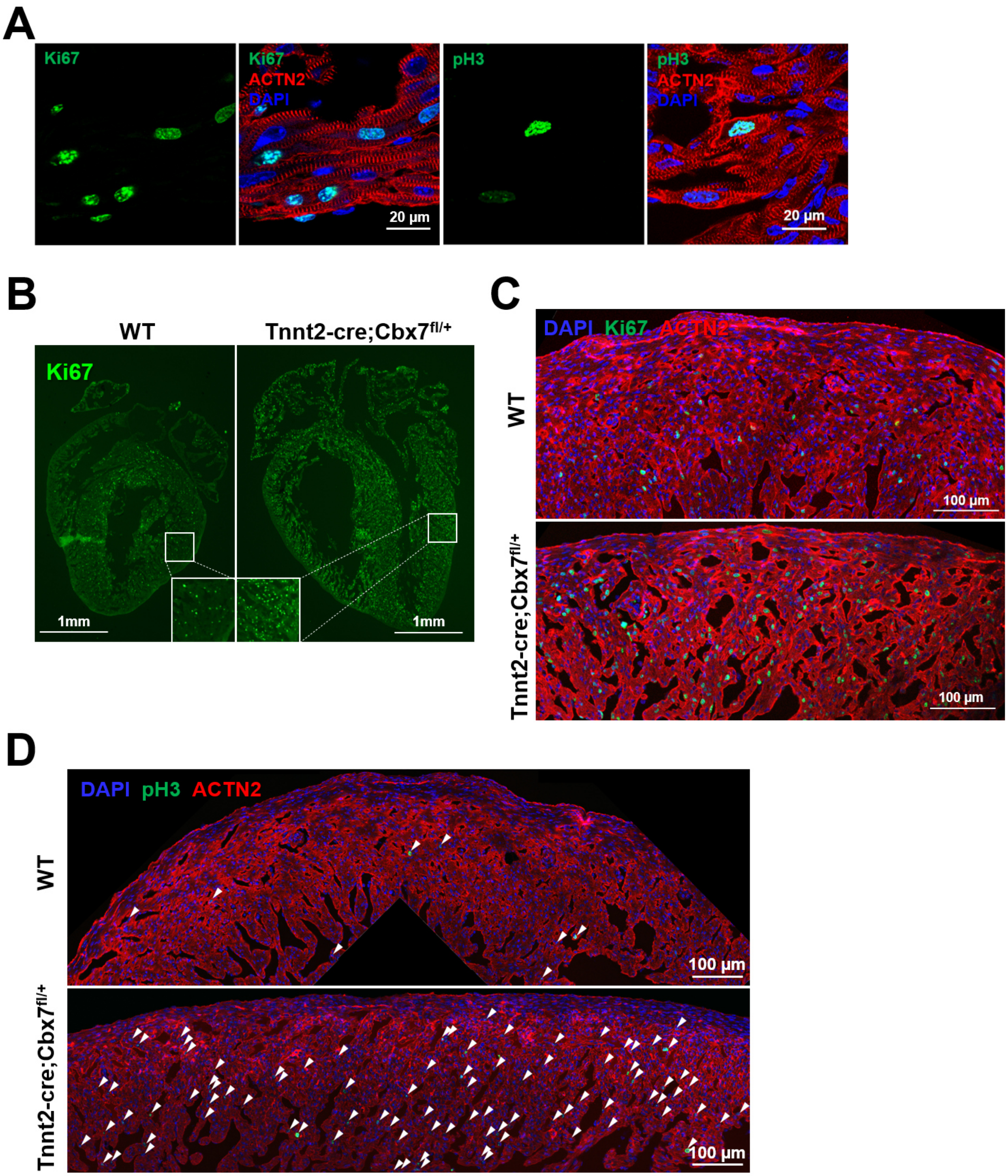


**Figure S5. Representative confocal microscopic images of neonatal mouse hearts immunostained for Ki67 and pH3. A.** immunohistochemistry for ACTN2 with either Ki67 (left) or pH3 (right) with WT neonatal (P0) mouse hearts. DAPI (blue). **B-D**. Low magnification images clearly showing an increased number of $Ki67^{+}$ CMs (**B-C**) and $pH3^{+}$ CMs (**D**) in Tnnt2-Cre;$Cbx7^{fl/+}$ mice compared to WT mice.

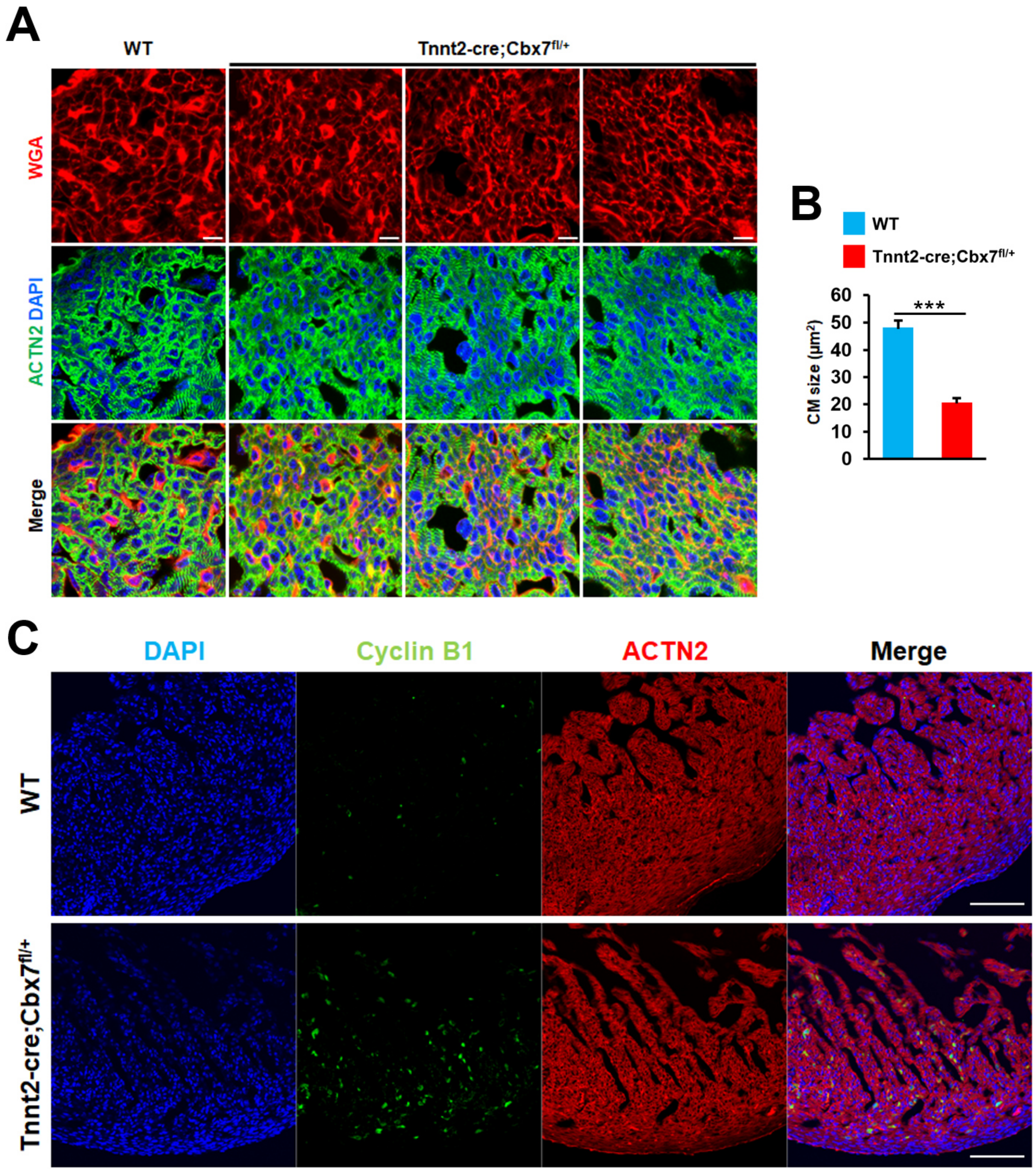


**Figure S6. Reduction in CM size and increase in number of Cyclin B$^+$ CMs in the neonatal heart by targeted inhibition of CBX7. A.** Representative confocal microscopic images for neonatal (P0) wild-type and Tnnt2-Cre;Cbx7$^{fl/+}$ mice stained for ACTN2 and WGA. Bar = 10 μm. **B.** Quantification of the CM size (double positive for ACTN2 and WGA). ***P < 0.001, Standard unpaired Student's t test, N = 3. More than 200 cells in each group were analyzed by using ImageJ software. **C.** Representative confocal microscopic images of the left ventricle of control and Tnnt2-Cre;Cbx7$^{fl/+}$ mice, immunostained for ACTN2 and Cyclin B1. DAPI (blue). Bar = 100 μm.

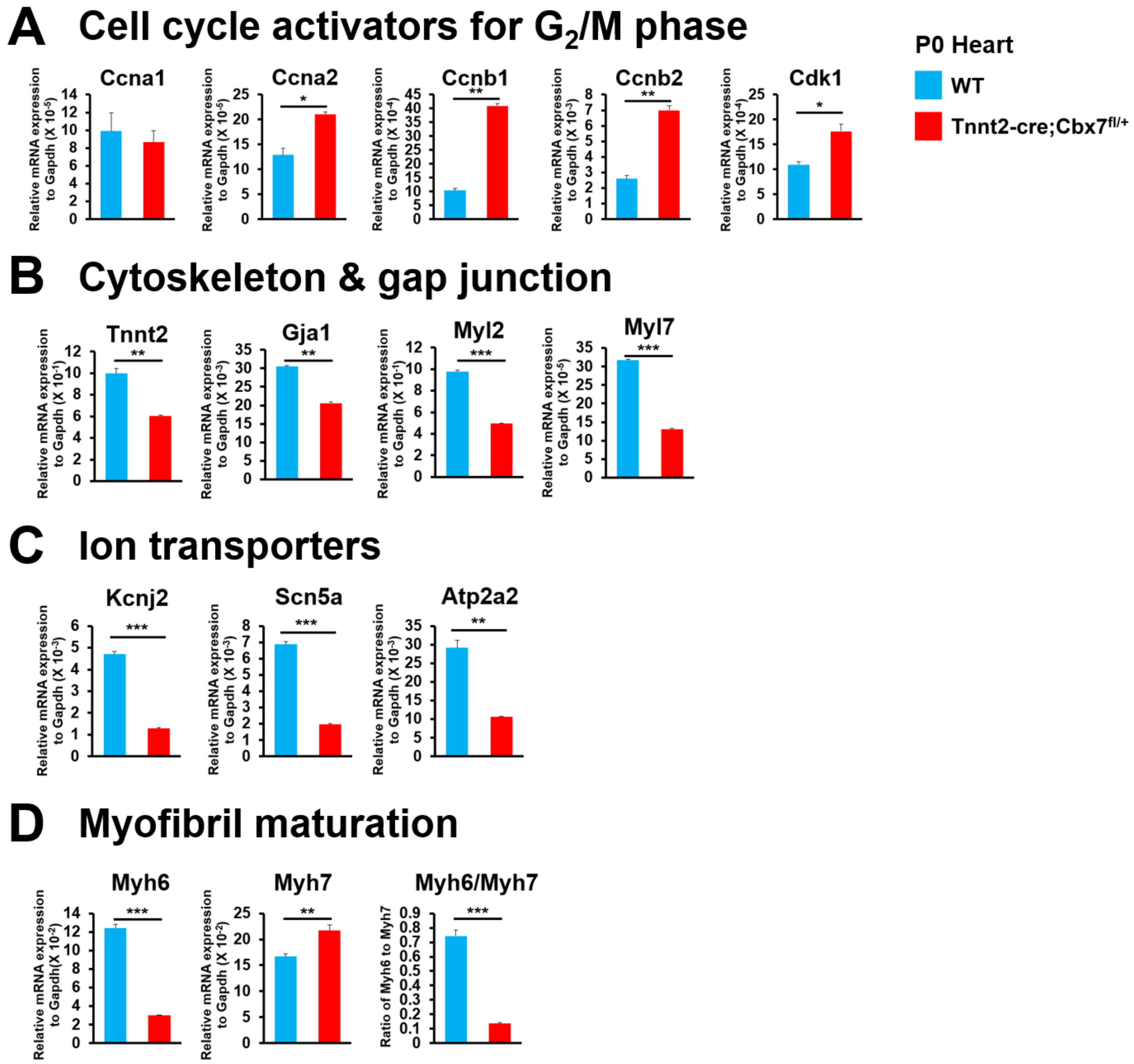


**Figure S7. Upregulation of mitotic signaling and downregulation of cardiac maturation-related genes by targeted inhibition of CBX7**. **A-E.** Representative qRT-PCR analyses for genes related to cell cycle activators (**A**), cytoskeletons & gap junctions (**B**), ion transporters (**C**), and myofibril maturation (**D**) with neonatal (P0) hearts derived from WT and Tnnt2-Cre;Cbx7$^{fl/+}$ mice. RNAs were isolated from whole hearts. Relative expression was normalized with Gapdh. *P < 0.05, **P < 0.01, ***P < 0.001. Standard unpaired Student's t test. N = 3, each with technical triplicates.

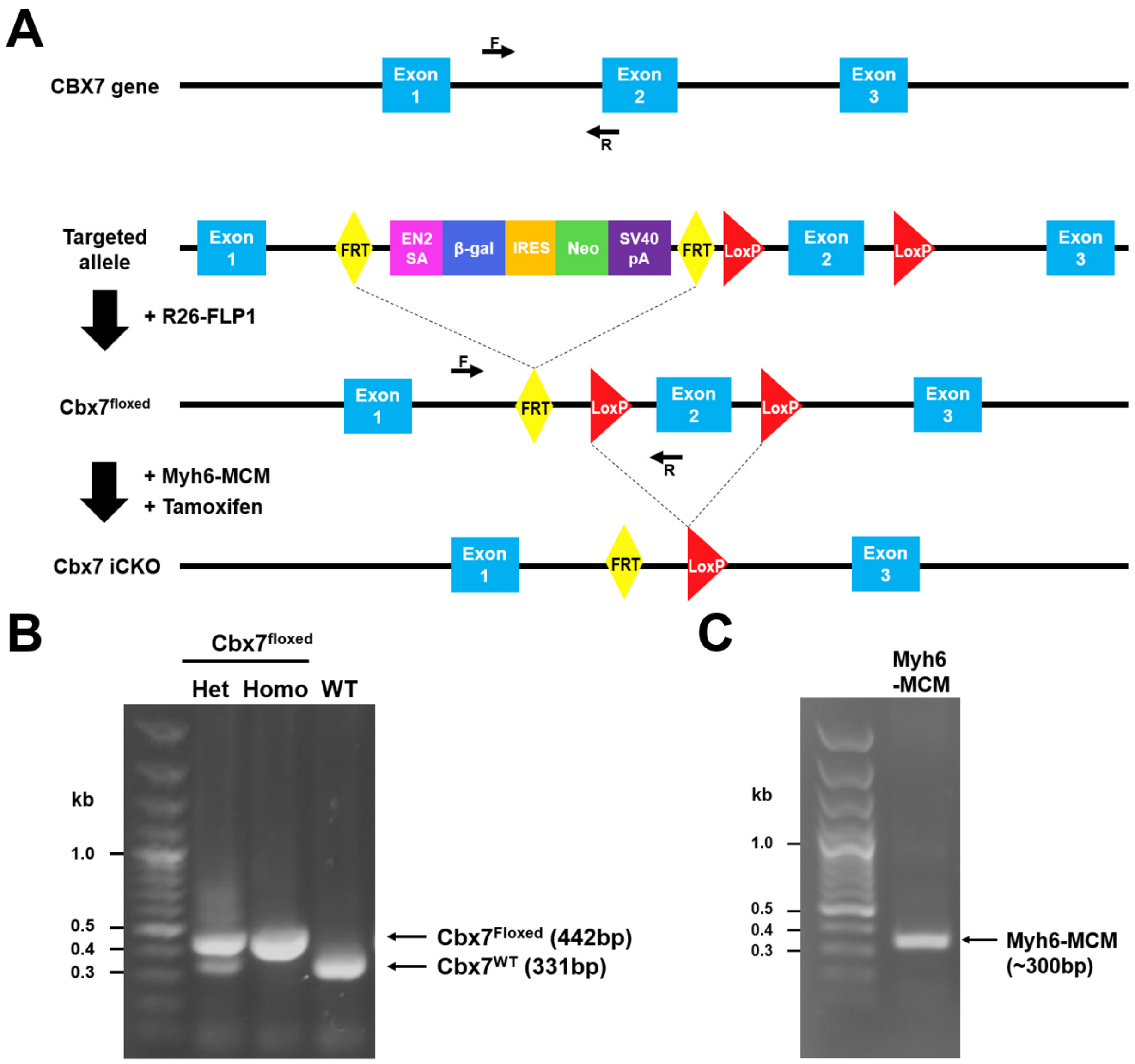


**Figure S8. A schematic and validation of the targeted Cbx7 allele for the tamoxifen inducible Cre-loxP system. A.** A schematic showing a Cbx7 allele and the targeted allele. The mice carrying the targeting vector (2nd from the top) were crossbred with R26-FLP1 mice leading to removal of the trapping cassette flanked by FRT (3rd from the top). The progeny were further crossbred with Myh6-MCM mice for Cre-mediated recombination to remove Exon 2 flanked by LoxP sites upon tamoxifen administration (Lowest), generating the inducible conditional knockout (iCKO). Arrows with “F” or “R” represent forward or reverse primers used for genotyping of the floxed Cbx7 allele in panel B. **B.** Representative PCR results using genomic DNA collected from the indicated mice detecting the floxed Cbx7 allele. **C.** Representative PCR results using genomic DNAs collected from the Myh6-MCM mice detecting the targeted Myh6 allele.

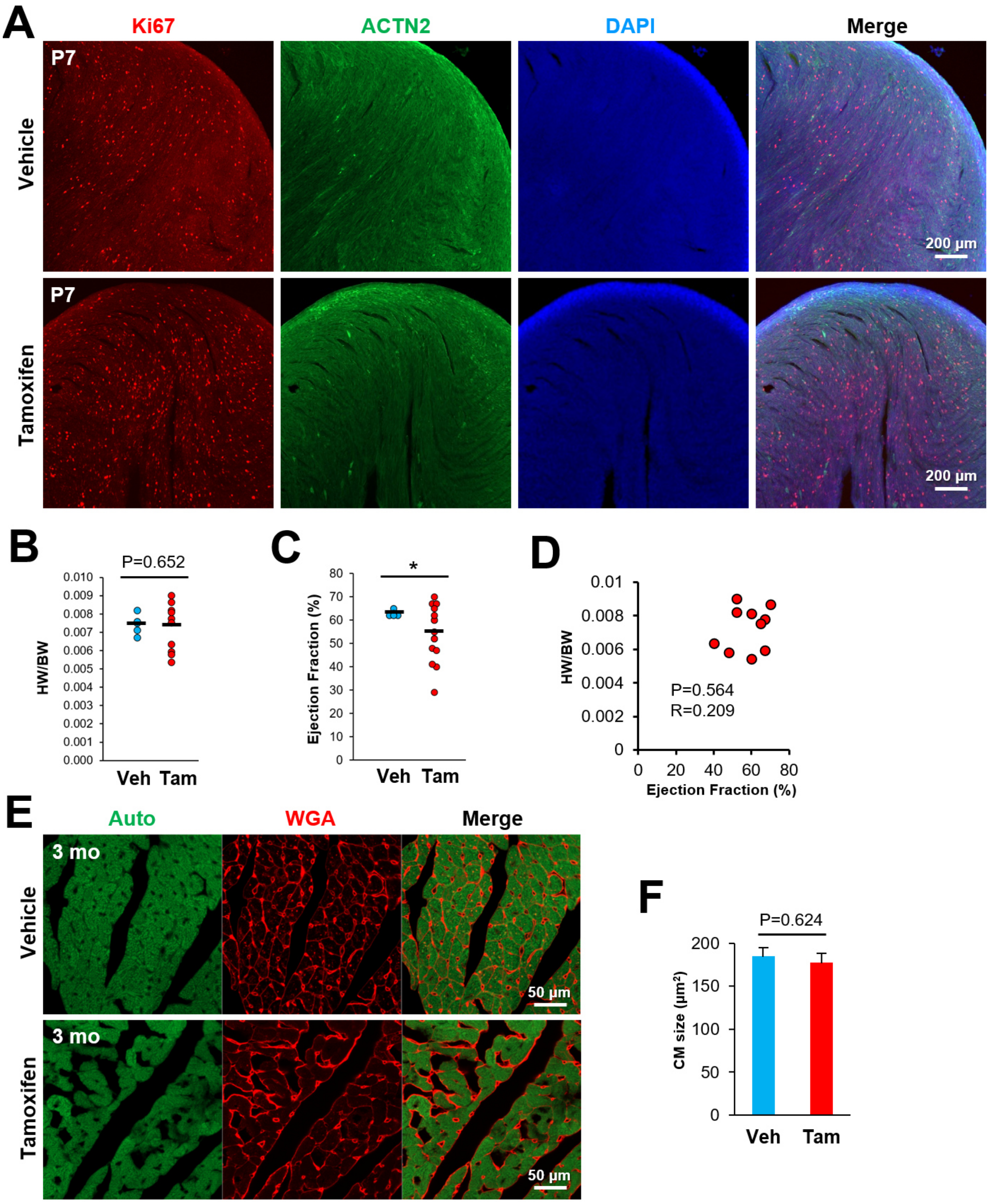


**Figure S9. Increased cardiomyocyte proliferation in the preadolescent heart and reduced cardiac function without changes in CM size in the adult heart by targeted inhibition of CBX7. A.** Representative confocal microscopic images of the left ventricle of Myh6-MCM;Cbx7$^{fl/fl}$

mice (P7), immunostained for Ki67 and ACTN2. DAPI (blue). Vehicle or tamoxifen were treated at P1. **B.** The heart to body weight ratio of Myh6-MCM;Cbx7$^{fl/fl}$ mice (3-month-old). Standard unpaired Student's t test, N = 5 to 10, each with technical triplicates. **C.** Echocardiography of Myh6-MCM;Cbx7$^{fl/fl}$ mice (3-month-old). *P < 0.05. Standard unpaired Student's t test. N = 5 to 13, each with technical triplicates. **D.** Correlation analysis between HW/BW and ejection fraction of Myh6-MCM;Cbx7$^{fl/fl}$ mice (3-month-old). R, Pearson's correlation coefficient. **E.** Representative confocal microscopic images of the left ventricle of Myh6-MCM;Cbx7$^{fl/fl}$ mice (3-month-old), stained with WGA. **F.** Measurement of the CM sizes. Standard unpaired Student's t test. More than 200 cells in each group were analyzed by using ImageJ software.

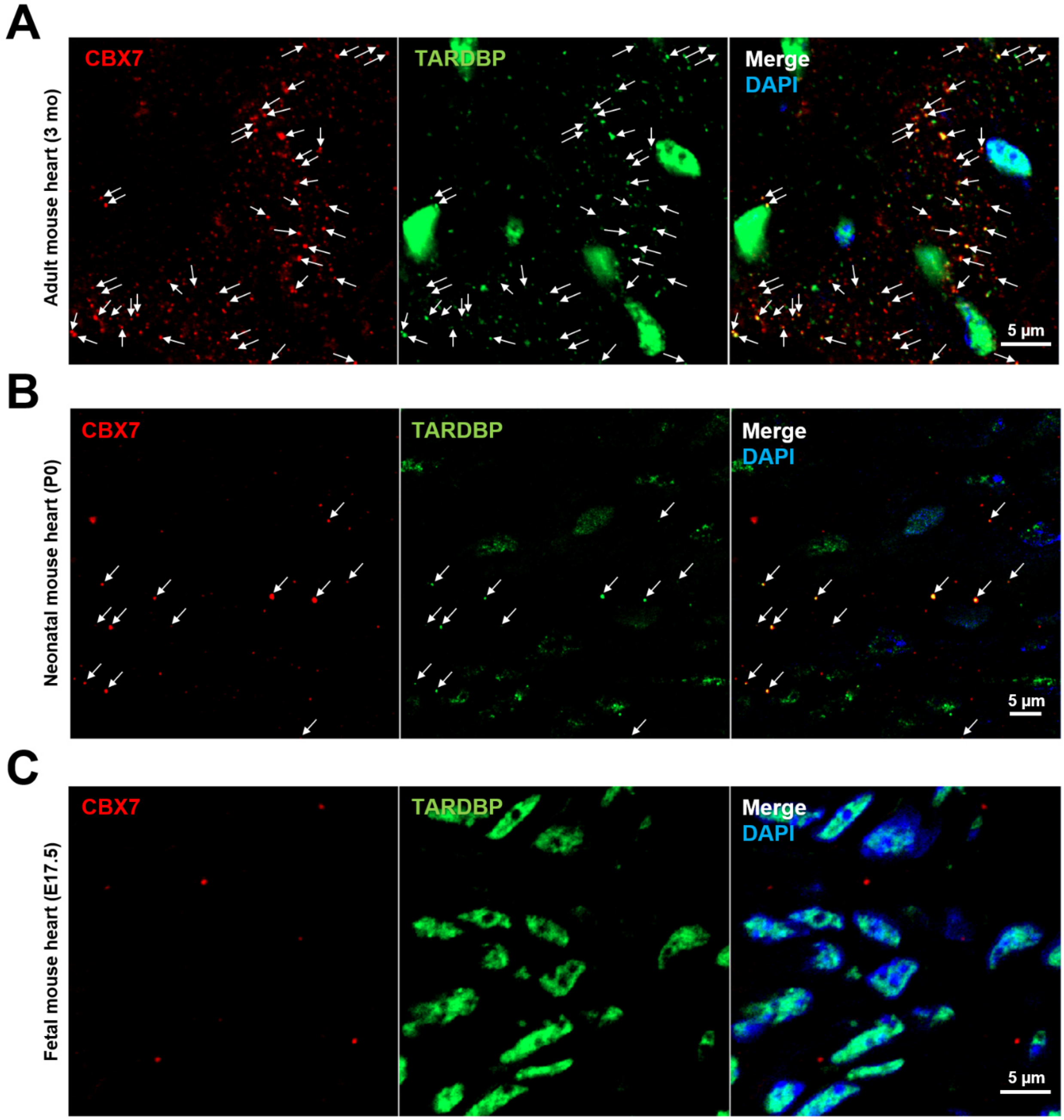


**Figure S10. CBX7 interacts with TARDBP in the postnatal heart. A-C.** Representative confocal microscopic images of adult (A), neonatal (B) and fetal (C) mouse hearts immunostained for CBX7 and TARDBP. DAPI (blue). Arrows indicate colocalization between CBX7 and TARDBP.

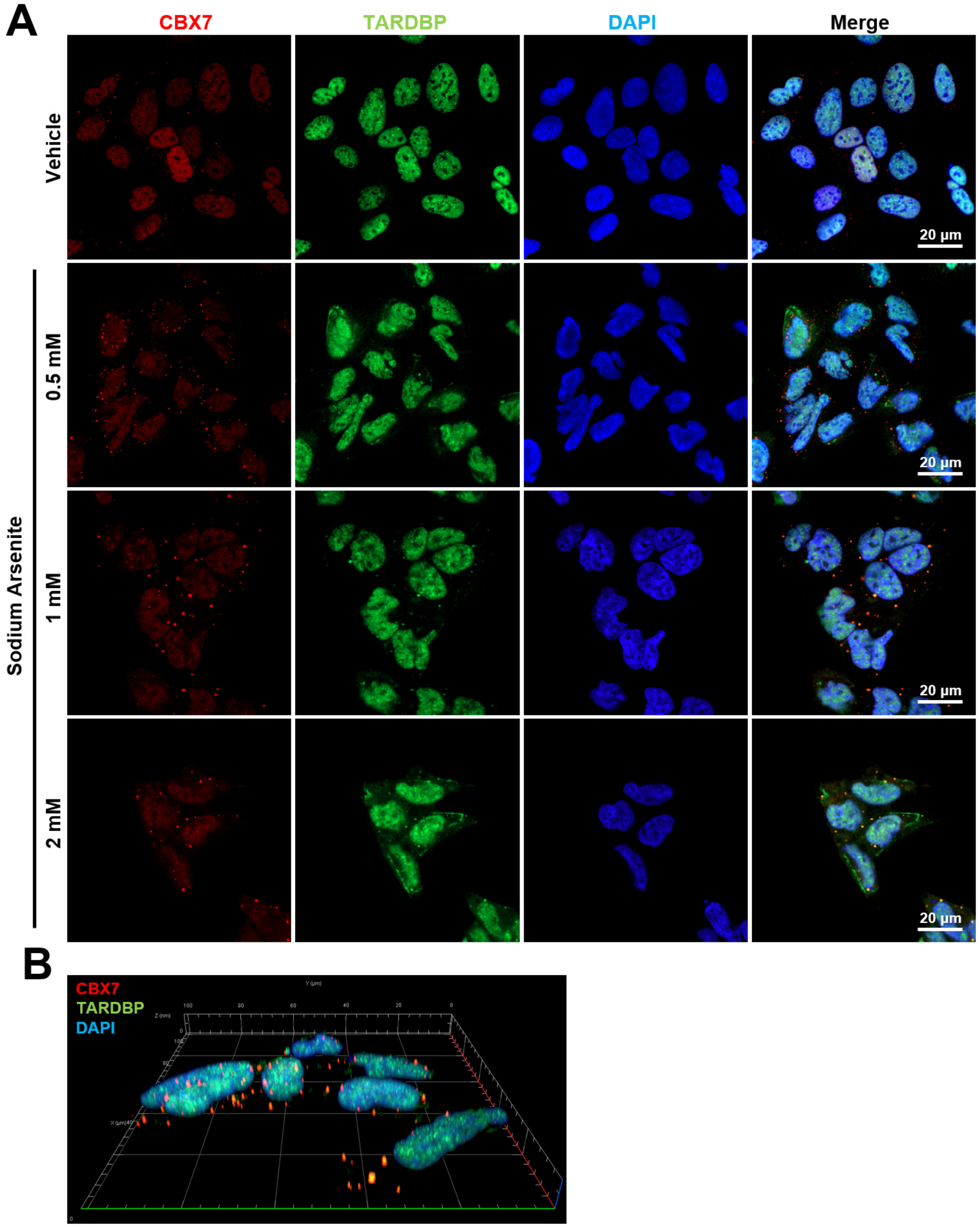


**Figure S11. Colocalization of CBX7 and TARDBP in HEK-293 cells in response to Sodium Arsenite. A.** Representative confocal microscopic images of HEK-293 cells immunostained for

CBX7 and TARDBP. DAPI (blue). Cells were treated with Sodium Arsenite at the indicated concentration for 1 hr. **B.** A representative three-dimensionally reconstructed image of Sodium Arsenite-treated HEK-293 cells (1 mM, 1 hr) immunostained for CBX7 and TARDBP. DAPI (blue).

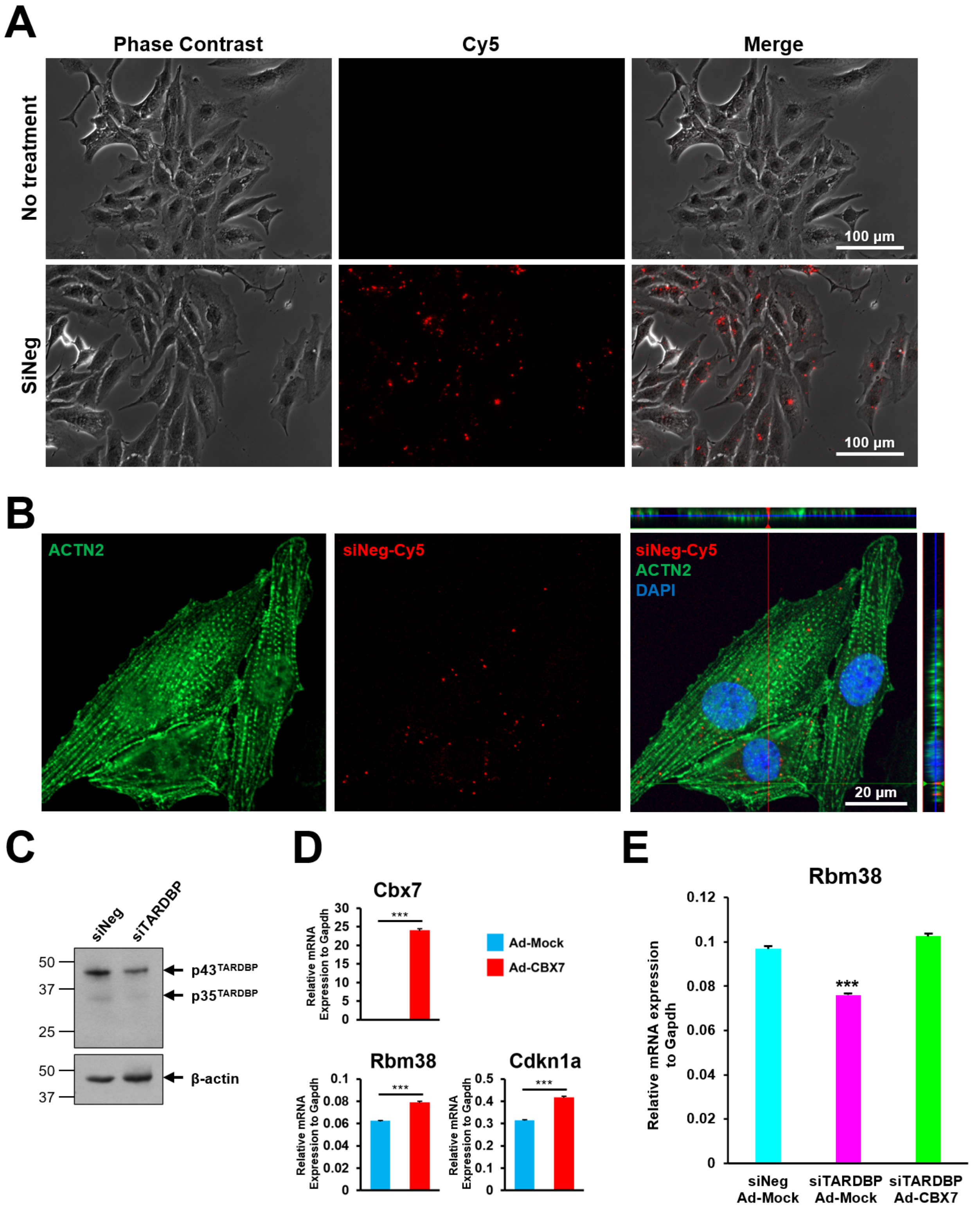


**Figure S12. Regulation of Rbm38 gene expression by TARDBP and CBX7 in HL-1 CMs. A.** Representative phase contrast and fluorescent microscopic images of HL-1 CMs 2 days after transfection of the negative siRNA control (siNeg) conjugated with fluorescent dye Cy5. **B.** Representative confocal microscopic images of HL-1 CMs immunostained for ACTN2 for validation of siRNA transfection. Cells were transfected with siNeg-Cy5 and subjected to immunocytochemistry 2 days later. DAPI (blue). Far right image is a representative orthogonal image of z-stacks. **C.** Validation of TARDBP knockdown in HL-1 CMs via western blotting. Cells were transfected with either siNeg-Cy5 or siTARDBP and subjected to western blotting 2 days later for TARDBP and β-actin. **D.** Validation of CBX7 overexpression in HL-1 CMs via qRT-PCR. Cells were treated with either Ad-Mock or Ad-CBX7 and subjected to qRT-PCR. Standard unpaired Student's t test. N = 3, each with technical triplicates. **E.** Positive regulation of Rbm38 by both CBX7 and TARDBP in HL-1 CMs. Cells were treated with indicated adenoviral particles and siRNAs. Two days later, cells were harvested and subjected to qRT-PCR for Rbm38 gene. one-way ANOVA test with Tukey HSD. N = 3, each with technical triplicates.

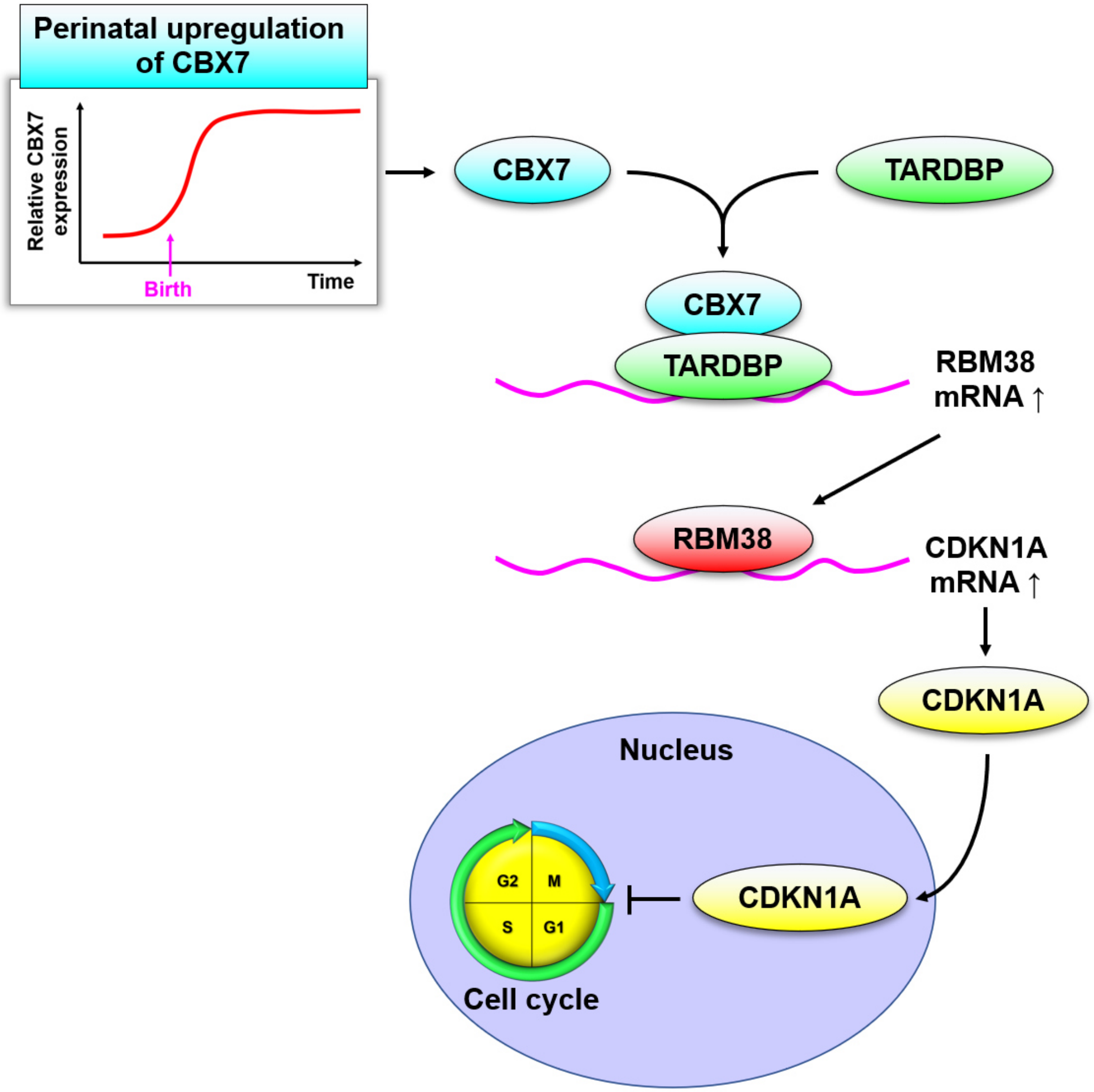


**Figure S13. A working model of how CBX7 represses CM proliferation.** In the postnatal heart, CBX7 interacts with TARDBP to increase RBM38 mRNA, a target of TARDBP. Translated RBM38 protein binds to CDKN1A transcripts, increasing CDKN1 mRNA. Then increased CDKN1A protein enters the nucleus and inhibits the cell cycle progression.

**Supplementary Table 1. Sequence of primers used in this study.**

**Supplementary Video 1. Wild type and Tnnt2-Cre;Cbx7$^{fl/+}$ mice at P1.**

**Supplementary Video 2. Death of Tnnt2-Cre;Cbx7$^{fl/+}$ mice at P1.**
